# Mechanisms regulating the frequency of inhibition-based gamma oscillations in primate prefrontal and parietal cortices

**DOI:** 10.1101/2022.04.26.489470

**Authors:** Guillermo Gonzalez-Burgos, Takeaki Miyamae, Nita Reddy, Sidney Dawkins, Chloe Chen, Avyi Hill, John Enwright, G Bard Ermentrout, David A. Lewis

## Abstract

In primates, the dorsolateral prefrontal (DLPFC) and posterior parietal (PPC) cortices are critical nodes in the network mediating cognitive functions including attention and working memory. Notably, during working memory tasks, gamma oscillations, usually prominent in layer 3 (L3), are induced in both DLPFC and PPC but exhibit higher frequency in DLPFC. These oscillation frequency differences might be crucial for working memory function, but the mechanisms producing different oscillation frequencies in monkey DLPFC and PPC remain poorly understood.

To investigate the basis of the DLPFC-PPC differences in oscillation frequency we studied GABA_A_R-mediated inhibition, which plays a crucial role in gamma oscillation production, in L3 pyramidal neurons (L3 PNs) from the rhesus monkey DLPFC or PPC. Recordings of GABA_A_R-mediated synaptic currents from L3 PNs, while suggesting a contribution to network synchronization in both areas, revealed no DLPFC-PPC differences in the strength or kinetics of GABA_A_R-mediated inhibition. Likewise, the expression of GABA_A_R genes in L3 PNs did not differ between regions.

In the absence of differences in inhibition, DLPFC L3 PNs showed greater dendritic spine density and higher expression of AMPAR and NMDAR subunit genes relative to PPC L3 PNs, suggesting that the excitatory synaptic drive onto L3 PNs could be stronger in the DLPFC. Simulations in computational models of the cortical microcircuit showed that, with constant synaptic inhibition, increasing the strength of recurrent excitatory synaptic drive increased the network oscillation frequency. Hence, the DLPFC-PPC differences in gamma oscillation frequency could depend on stronger recurrent excitation in the DLPFC relative to PPC.

**Significance statement:** Gamma oscillations may contribute to the neural substrate of working memory and exhibit a higher frequency in the prefrontal (DLPFC) than parietal (PPC) areas of primate cortex. To investigate the basis of these oscillation frequency differences which may be crucial for working memory encoding, we studied GABA_A_R-mediated inhibition on L3 pyramidal neurons (L3 PNs) from rhesus monkey DLPFC or PPC. Our data revealed no DLPFC-PPC differences in GABAAR-mediated inhibition but showed greater dendritic spine density in DLPFC L3 PNs, suggesting stronger excitatory synaptic drive. Simulations in computational network models showed that stronger recurrent excitatory synaptic drive increased the network oscillation frequency, suggesting that the higher oscillation frequency could depend on stronger recurrent excitation in the DLPFC relative to PPC.

## Introduction

Within the primate neocortex, the dorsolateral prefrontal (DLPFC) and posterior parietal (PPC) areas are key nodes in the network mediating visual working memory and attention (Goldman-Rakic, 1988; Christophel et al., 2017; Leavitt et al., 2017). The DLPFC and PPC are reciprocally connected by the principal axon projections from layer 3 pyramidal neurons (L3 PNs) (Selemon and Goldman-Rakic, 1988; Markov et al., 2014). These reciprocal connections likely contribute to the interactions observed between DLPFC and PPC during working memory tasks (Todd and Marois, 2005; Hart and Huk, 2020; Yu et al., 2020).

During the delay period of working memory tasks, the power of gamma frequency (30-100 Hz) oscillations increases in primate DLPFC and PPC (Pesaran et al., 2002; Howard et al., 2003; Roux et al., 2012; Wang et al., 2022). Therefore, gamma oscillations, which are predominantly localized to supragranular layers 2-3 (Bastos et al., 2018), might be an important neural substrate for working memory maintenance (Lundqvist et al., 2016; Miller et al., 2018). Working memory-related gamma oscillations exhibit higher frequency in monkey DLPFC relative to PPC, and are significantly correlated with spiking activity (Lundqvist et al., 2020). Hence, these higher-frequency oscillations in DLPFC might be crucial for encoding information maintained in working memory and for decoding the working memory-related information transmitted to other cortical areas (Lundqvist et al., 2020). Interestingly, a similar gradient from parietal to frontal areas of greater natural oscillation frequency has been reported in human cortex (Rosanova et al., 2009; Ferrarelli et al., 2012).

Despite the potential relevance of higher gamma oscillation frequency in DLPFC for working memory function (Lundqvist et al., 2020), the mechanisms that could generate these regional differences in oscillation frequency are poorly understood. Gamma oscillation production is thought to depend on rhythmic synaptic inhibition (Whittington et al., 2000; Buzsaki and Wang, 2012), which in the primate neocortex, as in rodents, seems to be produced by GABAergic interneurons (Banaie Boroujeni et al., 2021). Such rhythmic inhibition is elicited via GABA_A_ receptor-mediated inhibitory post-synaptic currents (GABA_A_R-IPSCs) that show properties critical for gamma oscillation production (Whittington et al., 2000; Buzsaki and Wang, 2012). For example, the oscillation frequency is higher if the GABA_A_R-IPSCs exhibit shorter duration and larger amplitude (Gonzalez-Burgos et al., 2015). Hence, the different oscillation frequencies in DLPFC and PPC might reflect different kinetics or strength of synaptic inhibition. However, little is known regarding the relative properties of GABA_A_R-mediated inhibition in these two areas of the primate neocortex.

To investigate the cellular basis of the DLPFC-PPC differences in oscillation frequency, we studied GABA_A_R-mediated inhibition by recording GABA_A_R-IPSCs from L3 PNs in acute slices from the DLPFC or PPC of rhesus monkeys. Our results suggest that GABA_A_R-IPSCs partake in L3 PN synchronization in both DLPFC and PPC but revealed no regional differences in the strength or kinetics of synaptic inhibition. Likewise, the expression of GABA_A_R subunit genes in L3 PNs did not show DLPFC-PPC differences.

Additionally, we found that, among L3 PNs showing no DLPFC-PPC differences in GABA_A_R-IPSCs, those from DLPFC had higher density of dendritic spines, the main site of excitatory input onto PNs (DeFelipe and Farinas, 1992). Moreover, the expression of AMPAR and NMDAR subunit genes was higher in L3 PNs from DLPFC relative to PPC. These data suggest that the excitatory synaptic drive onto L3 PNs could be stronger in DLPFC, hence promoting gamma oscillations with higher frequency (Brunel and Wang, 2003; Lundqvist et al., 2013).

Simulations of inhibition-based oscillations in computational models of the local cortical microcircuit showed that, while holding synaptic inhibition constant, progressively increasing the strength of AMPAR- and NMDAR-mediated drive from local recurrent connections progressively increased the network oscillation frequency. Hence, we suggest that the oscillation frequency differences between DLPFC and PPC during working memory tasks (Lundqvist et al., 2020) could depend on stronger recurrent excitation in the DLPFC relative to PPC.

## Materials and methods

### Animals

Housing and experimental procedures were conducted in accordance with USDA and NIH guidelines and were approved by the University of Pittsburgh Institutional Animal Care and Use Committee. For the electrophysiology experiments, tissue was obtained from 8 rhesus monkeys (*Macaca mulatta*, 7 males and 1 female; 27 to 47 months of age) that were experimentally naïve until entry into this study. These animals were used for experiments in a previous study of L3 PNs in DLPFC and PPC (González-Burgos et al., 2019). Tissue sections from 5 of the 8 monkeys were used for microarray analyses of gene expression in L3 PNs.

### Brain slice preparation

Slices were prepared as described previously (Gonzalez-Burgos et al., 2015; González-Burgos et al., 2019). Briefly, tissue blocks (Figure 1A) containing both banks of the principal sulcus (DLPFC area 46) or the intraparietal sulcus and adjacent lateral cortex (PPC areas LIP and 7a) were obtained after the animals were deeply anesthetized and perfused transcardially (Gonzalez-Burgos et al., 2015) with ice-cold sucrose-modified artificial cerebro-spinal fluid (sucrose-ACSF, in mM): sucrose 200, NaCl 15, KCl 1.9, Na_2_HPO_4_ 1.2, NaHCO_3_ 33, MgCl_2_ 6, CaCl_2_ 0.5, glucose 10 and kynurenic acid 2; pH 7.3–7.4 when bubbled with 95% O_2_-5% CO_2_. Slices were obtained from either DLPFC (4 male monkeys) or PPC (3 males, 1 female). Coronal plane slices (300 μm thick) were cut in a vibrating microtome (VT1000S, Leica Microsystems) while submerged in ice-cold sucrose-ACSF. Immediately after cutting, the slices were transferred to an incubation chamber at room temperature filled with the following ACSF (mM): NaCl 125, KCl 2.5, Na_2_HPO_4_ 1.25, glucose 10, NaHCO_3_ 25, MgCl_2_ 1 and CaCl_2_ 2, pH 7.3–7.4 when bubbled with 95% O_2_-5% CO_2_. Electrophysiological recordings were initiated 1 to 14 hours after tissue slicing was completed. All chemicals used to prepare solutions were purchased from Sigma Chemicals Company.

**Figure 1.**
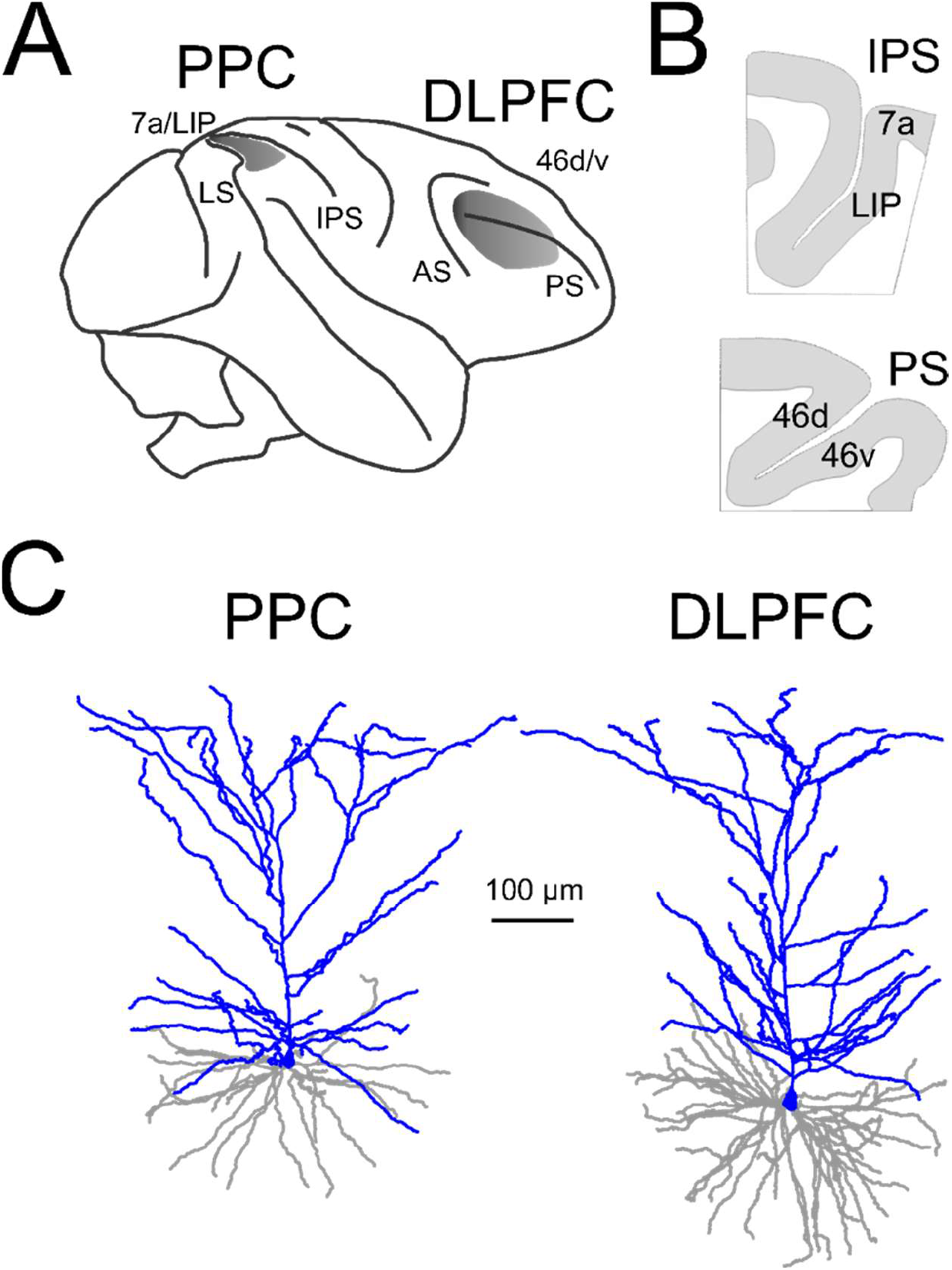
**A)** Cortical areas where tissue blocks were obtained for brain slice preparation. LS: lateral sulcus; IPS: intra-parietal sulcus; AS: arcuate sulcus; PS: principal sulcus. **B)** Schematic drawings representing the brain slices prepared from the PPC region surrounding the IPS (top), and the DLPFC surrounding the PS (bottom). Indicated are the areas where recordings from L3 PNs were obtained. **C)** Examples of the dendritic tree morphology of the recorded L3 PNs. Apical dendrites are shown in blue and basal dendrites in gray.

### Electrophysiological recordings

Slices were placed in a recording chamber superfused at 2-3 ml/min with the following ACSF (mM): NaCl 125; KCl 2.5; Na_2_HPO_4_ 1.25; NaHCO_3_ 25; glucose 10; CaCl_2_ 2; MgCl_2_ 1; CNQX 0.01; bubbled with 95 % O_2_ / 5 % CO_2_ at 30-32 °C. Whole-cell recordings were obtained from L3 PNs identified visually by infrared differential interference contrast video microscopy using Olympus BX51 or Zeiss Axioskop FS2 microscopes equipped with CCD video cameras (EXi Aqua, Q-Imaging). Recordings were obtained from L3 PNs located in either the medial or lateral bank of the principal sulcus containing DLPFC area 46, or in cytoarchitectonic areas LIP and 7a of the PPC, in the lateral bank of the intraparietal sulcus (Fig 1A). Recordings were obtained using Multiclamp 700A or 700B amplifiers (Axon Instruments) operating in voltage clamp mode, without employing series resistance compensation. Recordings were included in data analysis only if the resting membrane potential was ≤ −60 mV. All the neurons included in this study exhibited pyramidal morphology (Fig 1B). Recording pipettes had a resistance of 3-5 MΩ when filled with the following solution (mM): KGluconate 60; KCl 70; NaCl 10; EGTA 0.2; HEPES 10; MgATP 4; NaGTP 0.3, NaPhosphocreatine 14, biocytin 0.4 %, pH 7.2-7.3, adjusted with KOH. Biocytin was included to fill the cells during recordings, and study L3 PN morphology after the brain slices were fixed in paraformaldehyde and stained with the DAB method (González-Burgos et al., 2019). The series resistance was measured offline using Signal scripts written inhouse, as described previously (Miyamae et al., 2017). The series resistance did not differ (t_(56,1)_=1.495, p=0.141, Shapiro-Wilk Ln data, p=0.301) between the DLPFC (10.6±4.0 MΩ, n=28) and PPC (11.9±4.1 MΩ, n=30) samples of L3 PNs. Recorded sweeps were accepted for analysis only if the series resistance for a given sweep increased by less than 20% of the initial value.

GABA_A_R-mediated sIPSCs were analyzed using NeuroMatic software (Rothman and Silver, 2018), with a sliding threshold search algorithm (Kudoh and Taguchi, 2002). Using this approach, we detected ~ 200 events per L3 PN, and from the detected GABA_A_R-sIPSCs we measured: 1) the GABA_A_R-sIPSCs frequency, defined as the number of GABA_A_R-sIPSCs detected divided by the time window of recording that was analyzed; 2) the average GABA_A_R-sIPSC peak amplitude and the amplitudes of individual GABA_A_R-sIPSCs used to build the single-cell histograms of distribution of IPSC amplitudes; and 3) the weighted exponential decay time constant for the average GABA_A_R-sIPSCs waveform obtained for each L3 PNs, and for individual GABA_A_R-sIPSCs. The weighted exponential time constant is calculated as: Tau(w) = [(Tau(f) x A(f)) + (Tau(s) x A(s))]/(A(f)+A(s)); where Tau(w) is the weighted exponential time constant; Tau(f) and A(f), or Tau(s) and A(s), are the time constants and coefficients, of the faster (f) and slower (s) decay components. The time constants and coefficients were obtained after fitting a double exponential decay function to the decay phase of the GABA_A_R-sIPSCs,using the curve fitting routine in the NeuroMatic software tool (Rothman and Silver, 2018). The distributions of GABA_A_R-sIPSC Tau(w) were studied only if the curve fitting was successful for at least 100 of the individual events detected in each L3 PN. Given these criteria, the distribution of Tau(w) was studied in 26 of the 29 DLPFC L3 PNs, and 25 of the 32 PPC L3 PNs for which we had sIPSC-GABA_A_R data available.

### GABA and glutamate receptors subunit gene expression analysis

Previously published data from a microarray study of the transcriptomes of L3 PNs isolated from monkey DLPFC and PPC (González-Burgos et al., 2019) was interrogated for differential expression of GABA_A_R, AMPAR and NMDAR subunits. Briefly, individual L3 PNs were laser micro-dissected from coronal cryostat sections (12 μm thick) stained for Nissl substance as previously described (Datta et al., 2015). From each monkey, L3 PNs were collected from the dorsal and ventral banks of the principal sulcus (DLPFC area 46) or from the lateral bank of the intraparietal sulcus (PPC areas LIP and 7a) and pooled into samples of 100 cells. From each sample, RNA was extracted (RNeasy^®^ Plus Micro Kit, QIAGEN), subjected to a single round of amplification (Ovation Pico WTA system), labeled using the Encore Biotin module (Nugen) and loaded for transcriptome analysis on a GeneChip^™^ Rhesus Macaque Genome Array (ThermoFisher Scientific). Expression intensities were extracted from Affymetrix Expression Console using the RMA method (Irizarry et al., 2003) and transformed to log-scale (base 2), as previously described (González-Burgos et al., 2019). The analysis of GABA_A_R gene expression was limited to the six subunits (GABRA1, A2, A5; GABRB1, B2; GABRG2) that were highly expressed in the samples and that form most synaptic GABA_A_R complexes in the neocortex (Rudolph and Knoflach, 2011; Engin et al., 2018). The analysis of AMPAR and NMDAR gene expression included the genes (AMPAR: GRIA1, A2, A3, A4; NMDAR: GRIN1A, 2A, 2B) encoding the main subunits known to be present in excitatory synaptic sites (Hansen et al., 2021; Wichmann and Kuner, 2022). To assess regional differences in GABA, AMPA or NMDA receptors, we calculated composite scores for the relevant subunit genes in L3 PN samples from each animal as follows: 1) For each cortical region, Z scores for each transcript were determined for each of the 5 animals as Z= (x-m)/sd, where m and sd are the mean and standard deviation of the group of animals, and x is the expression ratio of each sample. 2) The mean Z score for all transcripts in each sample was then averaged across all 5 animals for each region and used as the composite score.

#### Histological processing and morphological reconstruction of biocytin-filled neurons

During recordings, L3 PNs were filled with 0.5% biocytin, and then visualized and reconstructed using procedures described previously (González-Burgos et al., 2019). Briefly, after recordings, the slices were immersed in 4% p-formaldehyde in 0.1 M phosphate-buffered saline (PBS) for 24-72 h at 4 °C. The slices were stored at −80 °C in a cryo-protection solution (33% glycerol, 33% ethylene glycol, in 0.1 M PBS) until processed. To visualize biocytin, the slices were re-sectioned at 60 μm, incubated with 1% H2O_2_, and immersed in blocking serum containing 0.5% Triton X-100 for 2–3 h at room temperature. The tissue was then rinsed and incubated with the avidin–biotin–peroxidase complex (1:100; Vector Laboratories) in PBS for 4 h at room temperature. Sections were rinsed, stained with the Nickel-enhanced 3,3’-diaminobenzidine chromogen, mounted on gelatin-coated glass slides, dehydrated, and cover slipped. Threedimensional reconstructions of the dendritic arbor were performed using the Neurolucida tracing system (MBF Bioscience). After tracing all basal dendrites of each L3 PN (Fig 7A), for each L3 PN a single primary basal dendrite was selected randomly and confirmed to have a true ending (tip of the basal dendrite not sectioned during slicing). On these dendrites, using differential interference contrast images obtained using 63X or 100X lenses, dendritic spines were identified throughout the entire length of each basal dendrite (Fig 7A). We calculated the mean spine density as the average of the spine number per micron for each basal dendrite. Identical conclusions regarding DLPFC-PPC differences in spine density were made in comparisons of the absolute number of dendritic spines per single basal dendrite, or when comparing estimates of the total number of spines per basal dendrite tree, obtained by multiplying the mean spine density by the total basal dendrite length for each L3 PN (data not shown).

### Statistics

Comparisons between group means were performed using Student’s t-test after the normality of distribution was assessed on the residuals of the data, or the Z-scores of the GABA_A_R subunit transcript data, using Shapiro-Wilk and D-Agostino Ksquare normality tests. When Shapiro Wilk or D-Agostino tests rejected the normality of distribution, we used natural logarithm transformation of the data, except when comparing Z-scores, which were not transformed. If normality of the distribution was rejected after log transformation, we used non-parametric tests, as indicated in each case. The statistical parameters of the histograms of distribution of IPSC properties were compared using non-parametric tests as well.

Most statistical comparisons using p value-based (frequentist) tests did not find significant differences between DLPFC and PPC. Hence, we combined the frequentist tests with Bayesian tests (Keysers et al., 2020) to obtain evidence for the absence of difference between the DLPFC and PPC groups. Using the JASP software (https://jasp-stats.org/), we computed the Bayes Factor (BF) for each comparison, thus estimating whether the data are better explained by the null hypothesis (H_0_: μ_DLPFC_ = μ_PPC_) or the alternative hypothesis (H_1_). Given that the higher gamma oscillation frequency in DLPFC than PPC (Lundqvist et al., 2020) would imply greater strength and/or faster decay kinetics of GABA_A_R-mediated inhibition (Gonzalez-Burgos et al., 2015), the alternative hypotheses were formalized as H1: μ_DLPFC_ > μ_PPC_ for the GABA_A_R-sIPSC amplitude, and H_1_: μ_DLPFC_ < μ_PPC_ for the GABA_A_R-sIPSC decay time constant comparisons, justifying one-tailed comparisons and computation of the BF_10_ (Keysers et al., 2020). For other comparisons, the alternative hypothesis was expressed as H_1_: μ_DLPFC_ ≠ μ_PPC_, and two-tailed tests were employed. As described previously (Keysers et al., 2020; van Doorn et al., 2021), Bayesian tests are considered to produce evidence favoring H_1_ when BF_10_ > 3, the evidence becoming stronger the larger BF_10_ is above 3. BF_10_ ≤ 1/3 is interpreted as evidence in favor of H_0_, and the smaller BF_10_ is below 1/3 the stronger the evidence for H_0_. Finally, 1/3 ≤ BF_10_ ≤ 3 suggests absence of evidence in favor of either H_1_ or H_0_. The outcome of the Bayes Factor analysis is reported in the figure legends.

### Computational modeling

The model consists of a network of 80 excitatory cells (e) and 20 inhibitory (i) spiking neurons. Each e and i cell is connected to every other e and i cell with a random strength drawn from a uniform distribution (0, 2) with mean 1, and divided by the number of inputs of that type. Each neuron obeys the following dynamics (Izhikevich, 2004):

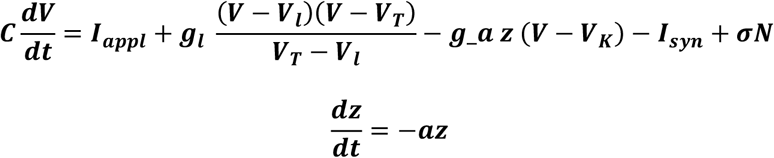

Where *z* represents adaptation a = 0.0125 (1/80ms), and *N* white noise (σ = 1) added to e and i cells. A random tonic drive *l_appl_*, 2.9 to 3.1 *μA/cm^2^* drives the e cells, and *l_appl_* = 0.1 to 0.4 *μA/cm^2^* is applied to the i cells. Each time *V(t)* hits *V_spike_, z* is incremented by *d* and *V* is reset to *V_r_.* For e cells, *g_l_* = 0.1, *V_l_* = −65, *V_T_* = −50, *V_r_* = −70, *d* = 1, *V_spike_* = 20, *V_K_* = −85. i cells are the same, but *V_r_* = −60 and *d* = 0 (no spike frequency adaptation). C = 1 *μF/cm^2^*, and all conductances are in *mS/cm^2^*, voltages are in mV. The conductance for the adaptation g_a was also varied, but typically has a value of 0.01.

Parameters are chosen so that at rest e cells and i cells have a membrane time constant of 10 ms. Synaptic currents have the form:

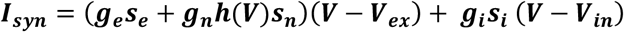

Where *V_in_*=-70, *V_ex_*=0 and

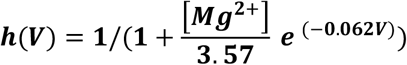

Every neuron to which a given cell is connected to, contributes its own synaptic current. The synaptic gating variables satisfy:

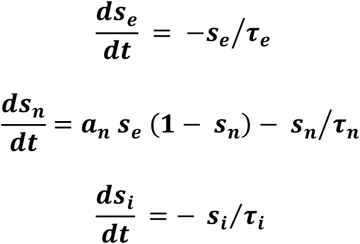

Here *s_e_*, *s_n_* and *s_i_*, are AMPA, NMDA and GABA synapses, respectively. The model for NMDA includes Mg^2+^ block with Mg^2+^ set to 0.5 mM (very similar effects of changing the recurrent excitatory conductance were obtained with simulations using Mg^2+^=1.0 mM). Each time an e cell fires, *s_e_* is incremented by 1; each time an i cell fires, its corresponding *s_i_* is incremented by 1. For e-e connections, G_*ee*_ and G_*ne*_ were independently varied between 0.1 and 1.0 mS/cm^2^. For e-i connections, G_*ni*_ = was held constant at 0.1 mS/cm^2^, and for i-e and i-i connections, G_*ie*_ = G_*ii*_ = 0.5 mS/cm^2^ were held constant. The exponential decay time constants of each synaptic conductance were, *τ_ee_* = *τ_ei_* = 1 ms, *τ_ne_* = *τ_ni_* = 80 ms, *τ_ie_* = *τ_ii_* = 4 ms. All simulations were performed using XPPAUT; the code is available from the authors. Euler’s method was used with a step size of 0.05 msec. Power spectra were taken on the *s_e_*, the e-e synaptic gating variable, which is the sum of all the excitatory currents into the excitatory cells. Reported are the peak oscillation frequency and peak oscillation power for each individual network with a particular combination of G_*ee*_ and G_*ne*_ values.

## Results

### GABA_A_R-mediated synaptic inhibition exhibits similar strength in DLPFC and PPC L3 PNs

We first determined whether GABA_A_R-mediated synaptic inhibition is stronger in L3 PNs from DLPFC relative to PPC, since stronger inhibition may generate higher gamma oscillation frequency (Gonzalez-Burgos et al., 2015) and hence could contribute to the higher oscillation frequency in DLPFC (Lundqvist et al., 2020). To test this idea, we recorded GABA_A_R-mediated spontaneous IPSCs (GABA_A_R-sIPSCs) from L3 PNs in acute slices from DLPFC area 46 (n=29 L3 PNs), or PPC areas 7a/LIP (n=32 L3 PNs), as illustrated in Fig 1.

As reported previously for DLPFC (Gonzalez-Burgos et al., 2015; Medalla et al., 2017), L3 PNs in both regions exhibited relatively frequent GABA_A_R-sIPSCs (Fig 2A), with substantial cell-to-cell variability in the GABA_A_R-sIPSC frequency (Fig 2B). The mean GABA_A_R-sIPSC frequency did not differ between DLPFC and PPC L3 PNs, whereas Bayesian analysis supported the absence of a difference in GABA_A_R-sIPSC frequency between cortical areas (Fig 2B). We next compared the strength of GABA_A_R-mediated inhibition in L3 PNs from DLPFC and PPC, by measuring the peak amplitude of the GABA_A_R-sIPSCs. The mean GABA_A_R-sIPSC amplitude (Fig 2C) did not differ between DLPFC and PPC L3 PNs, and Bayesian analysis supported no difference in GABA_A_R-sIPSC amplitude between regions (Fig 2D).

**Figure 2.**
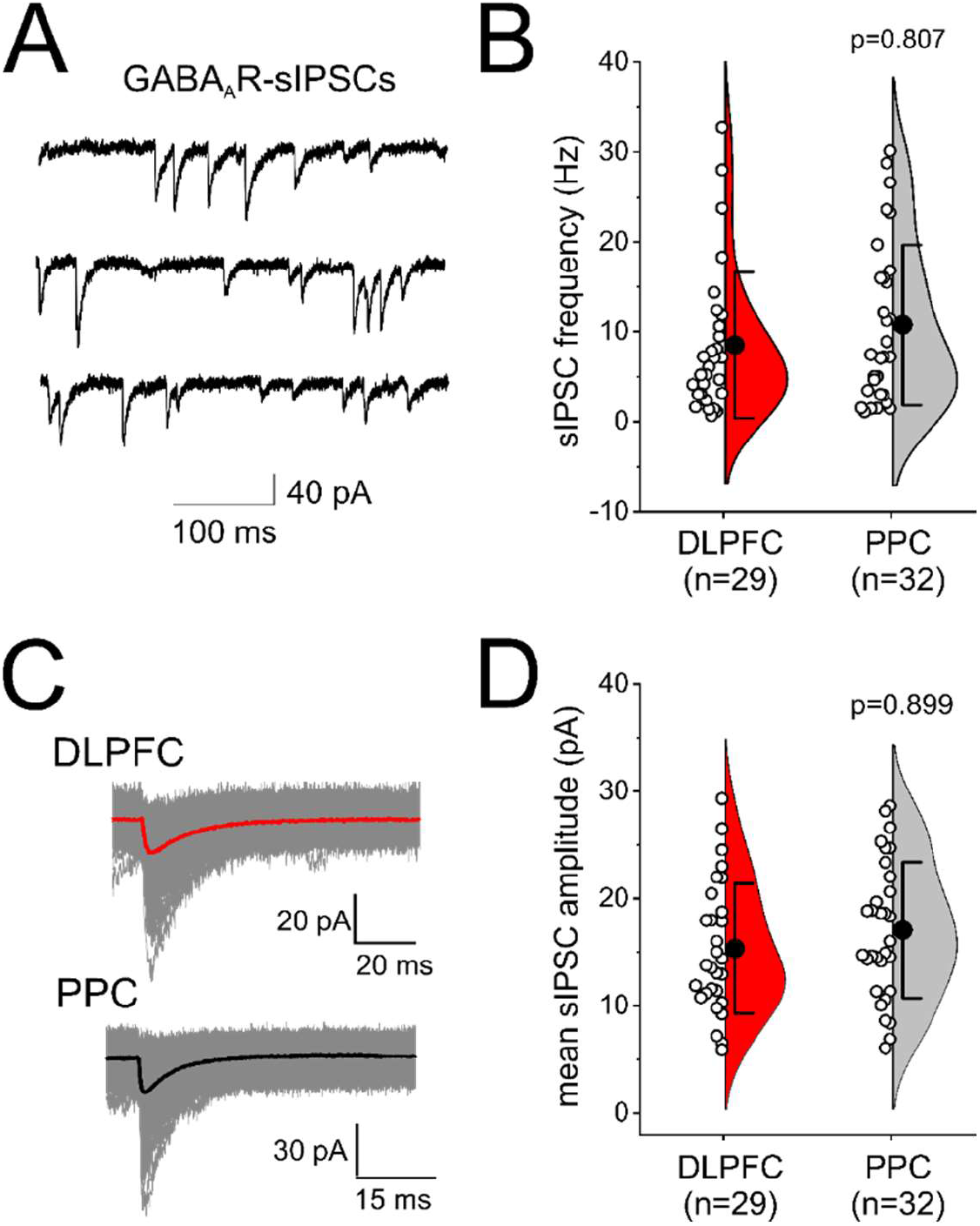
Frequency and amplitude of GABA_A_R-sIPSCs in DLPFC and PPC L3 PNs. **A)** Examples of GABA_A_R-sIPSCs. **B)** Violin plots of the GABA_A_R-sIPSC frequency. Here and in all other figures, the violin plots display a smoothed Kernel density plot, together with the data for individual cells (circles), the 25-75% interquartile range (bar) and the mean±SD. Student’s t-test and Bayes Factor analysis (BF_10_=0.157, median posterior d=-0.107, 95% CI: [-0.420, −0.004]), were performed after Ln transformation (Shapiro Wilk test, p= 0.812). **C)** Superimposed individual GABA_A_R-sIPSCs (gray traces) and average waveforms (black or red traces). **D)** Violin plot of the GABA_A_R-sIPSC peak amplitudes. The statistical comparisons were performed after Ln transformation (Shapiro Wilk p= 0.548). Bayes Factor analysis, BF_10_=0.126, median posterior d=-0.089, 95% CI: [−0.369, −0.004].

Consistent with the comparisons of mean GABA_A_R-sIPSC amplitude (Fig 2C,D), L3 PNs from DLPFC and PPC had very similar single-cell distributions of GABA_A_R-sIPSC amplitudes (Fig 3A). Statistical comparisons did not reveal differences between regions in these distributions (Fig 3B), showing that GABA_A_R-mediated inhibition is not stronger in L3 PNs from DLPFC. Moreover, parameters assessing the shape of the single-cell distributions (coefficient of variation, mode and skewness) did not differ between areas (Fig 3C), although Bayesian analyses did not provide strong evidence for the absence of difference in these parameters.

**Figure 3.**
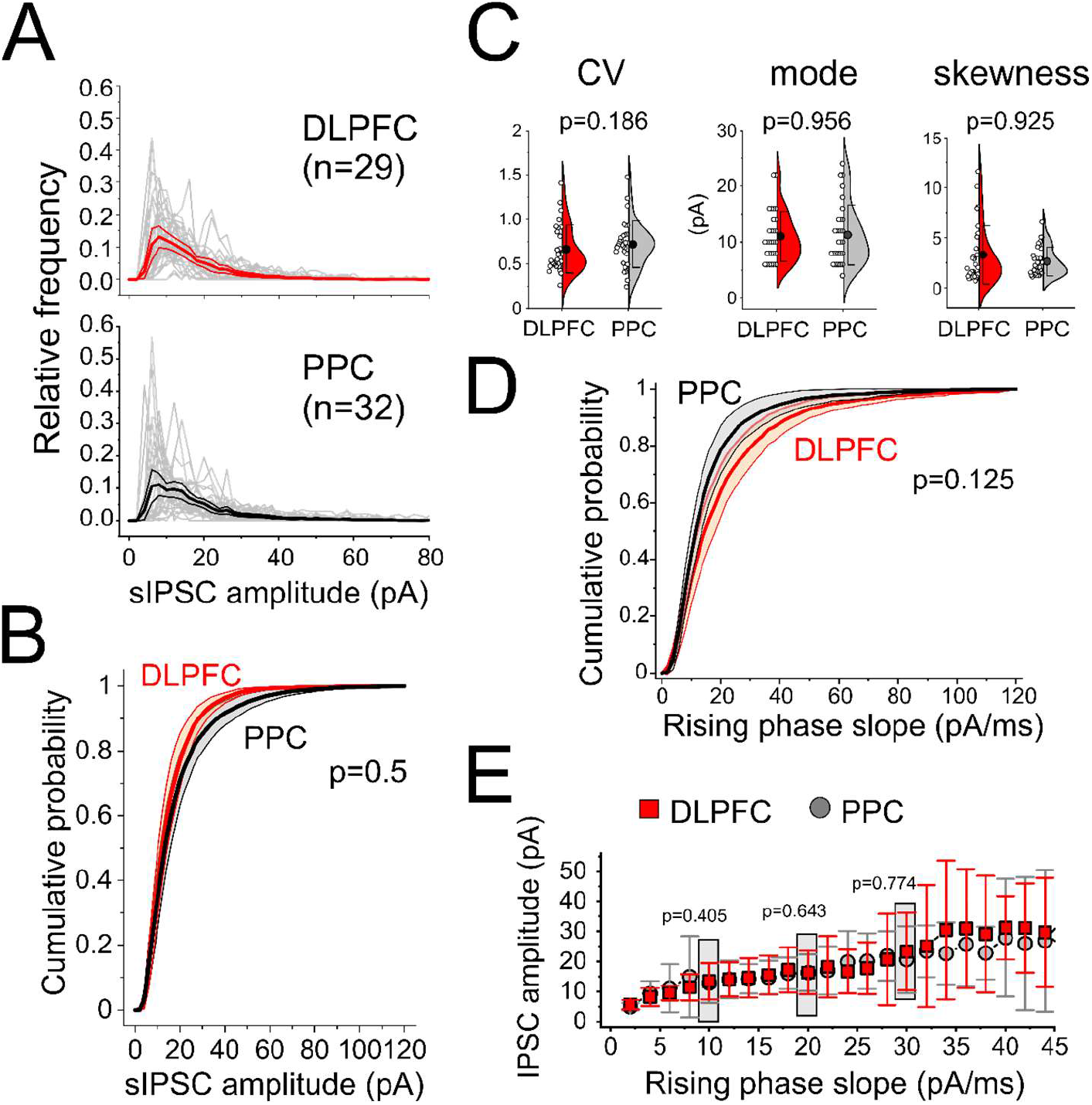
Distributions of GABA_A_R-sIPSC amplitudes in DLPFC and PPC L3 PNs. **A)** Histograms of distribution of GABA_A_R-sIPSC amplitude for single L3 PNs (gray lines) and the average (red and black lines) together with the 95% confidence intervals. **B)** Average cumulative probability distributions of the GABA_A_R-sIPSC amplitude in DLPFC and PPC L3 PNs, together with the 95% confidence intervals. Comparisons with Kolmogorov-Smirnov test. **C)** Violin plots for the statistical parameters of the distributions of GABA_A_R-sIPSC amplitudes. The circles are the parameters computed in the distribution of single cells. Shown are the p values of Mann-Whitney U test comparisons. Bayes Factor analysis, CV: BF_10_=0.434, median posterior d=0.231, CI: [-0.229,0.714]; mode: BF_10_=0.267, median posterior d=0.006, 95% CI: [-0.481, 0.466]; skewness: BF_10_=0.274, median posterior d=0.016, 95% CI: [-0.463, 0.495]. **D)** Average cumulative probability distributions of the slopes of the rising phase of the GABA_A_R-sIPSCs.Comparisons with Kolmogorov-Smirnov test. **E)** Plot of peak amplitude versus slope of the rising phase of the GABA_A_R-sIPSC. The GABA_A_R-sIPSC amplitude in the 10, 20 or 30 pA/ms slope groups was compared using Mann-Whitney U tests (Shapiro Wilk tests, 10 pA/ms, p= 9.1×10^-7^; 30 pA/ms, p=4.1×10^-5^; Ln-transformed 20 pA/ms, p= 0.012). Bayes Factor analysis, 10 pA/ms: BF_10_=0.365, median posterior d=0.190, CI: [-0.264,0.662]; 20 pA/ms: BF_10_=0.270, median posterior d=-0.049, 95% CI: [-0.518, 0.415]; 30 pA/ms: BF_10_=0.296, median posterior d=0.022, 95% CI: [-0.500, 0.547].

The strength of GABA_A_R-mediated inhibition onto L3 PNs could differ between DLPFC and PPC specifically for perisomatic GABA synapses, believed to play a crucial role in gamma oscillation production (Whittington et al., 2000; Buzsaki and Wang, 2012). GABA_A_R-IPSCs elicited by perisomatic synapses display faster slope of the rising phase and larger amplitudes than those elicited at more distal synapses (Xiang et al., 2002; Ali and Thomson, 2008; Donato et al., 2021). Thus, to compare synaptic currents likely originating from more proximal or more distal synapses, we sorted the GABA_A_R-sIPSCs using the slope of their rising phase and compared between DLPFC and PPC L3 PNs the GABA_A_R-sIPSC amplitudes across the range of rising phase slopes. The distributions of rising phase slopes largely overlapped between DLPFC and PPC L3 PNs (>90% of the GABA_A_R-sIPSCs showed slopes ≤45 pA/ms in both DLPFC and PPC L3 PNs) and did not differ between areas (Fig 3D). As expected for IPSCs originated perisomatically, the GABA_A_R-sIPSCs with faster rising phase had larger amplitudes, whereas IPSCs with slower rising phase were smaller, as expected for IPSCs elicited more distally (Fig 3E). Across the range of rising phase slopes, the GABA_A_R-sIPSC amplitude did not differ between DLPFC and PPC L3 PNs (Fig 3E), suggesting that GABA synapses have comparable strength in DLPFC and PPC L3 PNs irrespective of a perisomatic or distal location.

### GABA_A_R-mediated synaptic inhibition exhibits similar decay kinetics in DLPFC and PPC L3 PNs

Although our analysis of GABA_A_R-sIPSC amplitude did not reveal any DLPFC-PPC differences (Fig 2,3), our prior computational study showed that even with constant GABA synapse strength, the decay kinetics of the GABA_A_R-sIPSC significantly modulates gamma oscillation frequency (Gonzalez-Burgos et al., 2015). Hence, we assessed the decay kinetics of the GABA_A_R-sIPSCs by measuring their exponential decay time constant (Fig 4A). We found that the average decay time constant was similar in L3 PNs from DLPFC and PPC (Fig 4B). Moreover, the decay time constants showed wide distributions in both DLPFC and PPC L3 PNs (Fig 4C), and the distributions did not differ between areas (Fig 4D). The statistical parameters that measure the shape of the distributions were similar between areas, although the distributions exhibited greater CV and skewness in PPC (Fig 4E).

**Figure 4.**
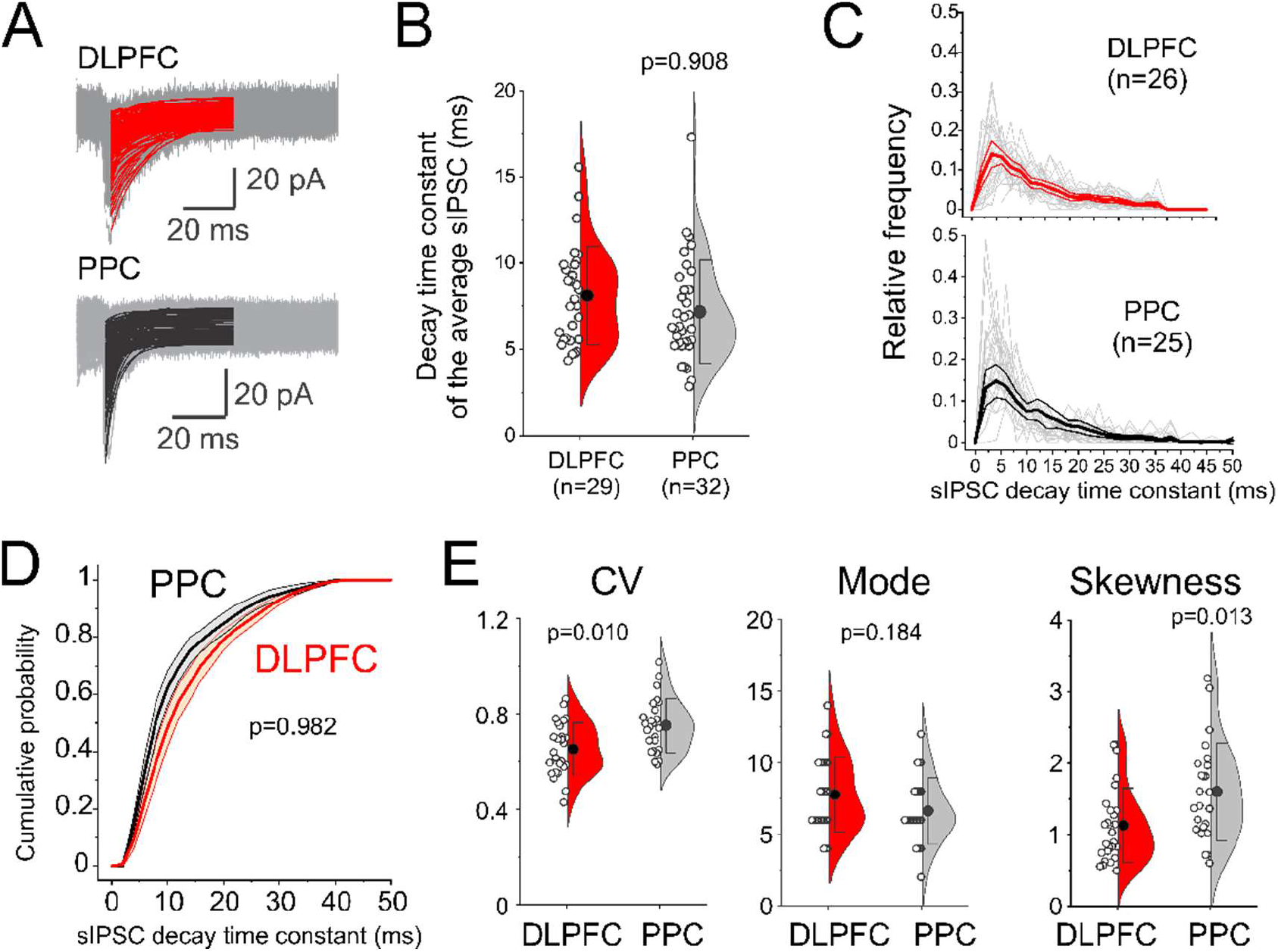
Legend. Decay time constants of the GABA_A_R-sIPSCs recorded from DLPFC and PPC L3 PNs. **A)** Superimposed individual GABA_A_R-sIPSCs (gray traces) and the double exponential curves fitted to the decay phase of the individual synaptic currents (red and black traces). **B)** Plots of the GABA_A_R-sIPSC decay time constant. Comparisons using Student’s t-test and Bayes Factor analysis performed after Ln transformation (Shapiro Wilk test, p= 0.869). Bayes Factor analysis: BF_10_=0.114, median posterior d=0.081, 95% CI: [−0.003, 0.347]. **C)** Histograms of distribution of GABA_A_R-sIPSC decay time constant for single L3 PNs (gray lines) and the average distributions (red and black lines) together with the 95% confidence intervals. The distributions were studied only for 26 and 25 DLPFC and PPC L3 PNs with adequate curve fitting for at least 100 events (see Materials and Methods). **D)** Average cumulative probability distributions of the GABA_A_R-sIPSC decay time constant in DLPFC and PPC L3 PNs, together with the 95% confidence intervals. Comparisons with Kolmogorov-Smirnov test. **E)** Plots of the statistical parameters of the distributions of GABA_A_R-sIPSC decay time constants in single L3 PNs. Circles indicate the parameters computed in the distribution of each cell. Comparisons with Mann-Whitney U tests. Bayes Factor analysis, CV: BF_10_=5.287, median posterior d=0.662, CI: [0.101,1.238]; mode: BF_10_=0.583, median posterior d=-0.313, 95% CI: [-0.860, 0.189]; Skewness: BF_10_=3.609, median posterior d=0.606, 95% CI: [0.061, 1.188].

The decay kinetics of the GABA_A_R-sIPSCs could differ between DLPFC and PPC exclusively in the subsets of GABA_A_R-sIPSCs directly involved in synchronizing networks of L3 PNs, which cannot be detected in the analysis of GABA_A_R-sIPSCs in single cells. Hence, to study specifically GABA_A_R-sIPSCs involved in synchronization, we recorded simultaneously from pairs of adjacent L3 PNs (Fig 5) and searched for GABA_A_R-sIPSCs that were synchronized between the L3 PNs in a pair.

**Figure 5.**
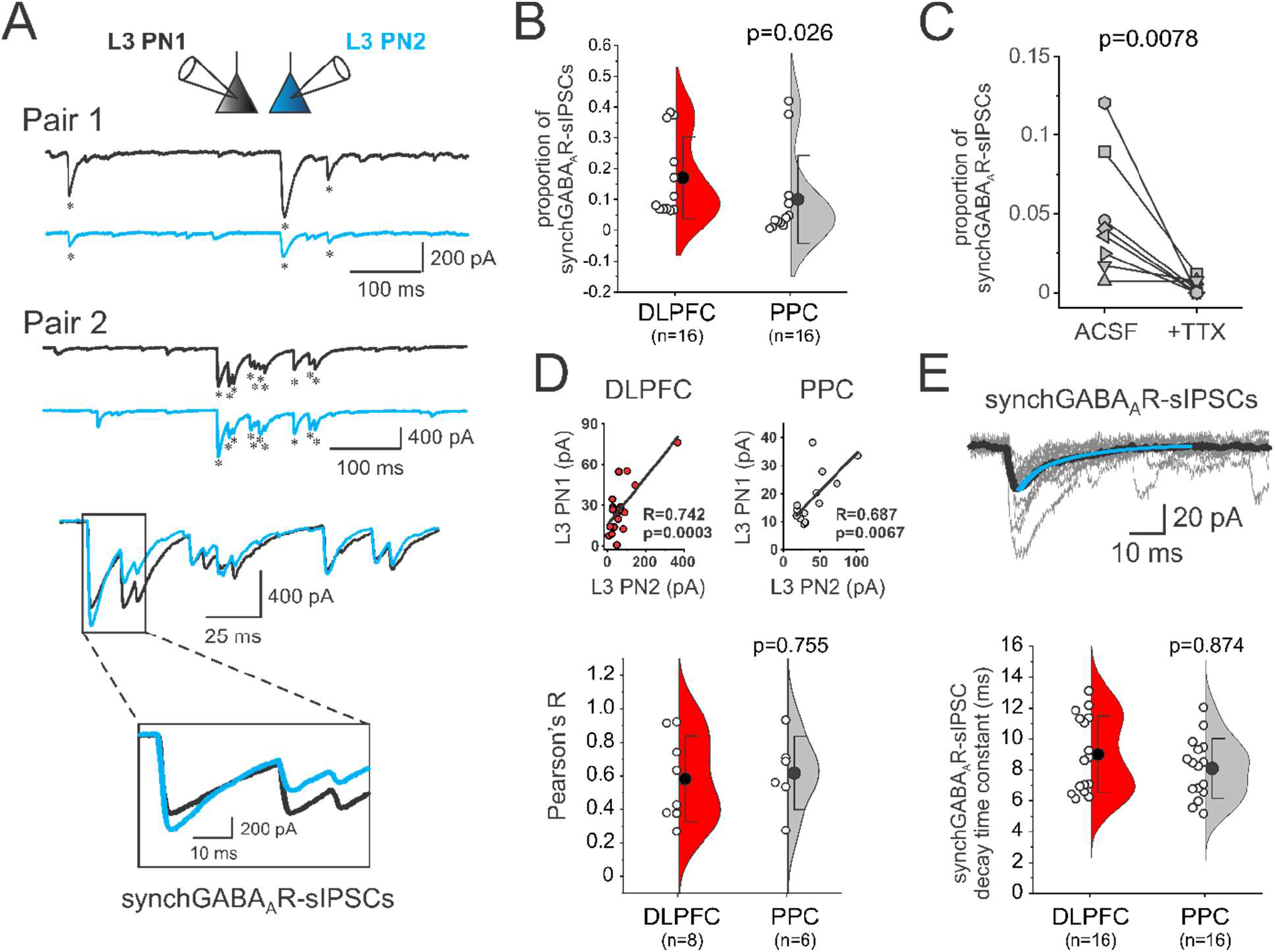
Legend. Synchronized GABA_A_R-sIPSCs in simultaneously recorded pairs of DLPFC and PPC L3 PNs. **A)** Synchronized GABA_A_R-sIPSCs (synchGABA_A_R-sIPSCs) recorded from two example pairs of L3 PNs in DLPFC (Pair 1 and Pair 2, respectively). Asterisks indicate GABA_A_R-sIPSCs designated as synchGABA_A_R-sIPSCs. The bottom two panels show at expanded time scales the burst of synchGABA_A_R-sIPSCs highlighted in Pair 2. **B)** Violin plots of the proportions of synchGABA_A_R-sIPSCs out of total GABA_A_R-sIPSCs in DLPFC and PPC L3 PNs. Shown are the results of statistical comparisons performed after Ln transformation of the data (Shapiro Wilk test, Ln-transformed data p=0.311). Bayes Factor analysis: BF_10_=1.148, median posterior d=0.557, 95% CI: [-0.178, 1.406]. **C)** Inhibition of action potential firing with tetrodotoxin (1 μM) abolishes the synchGABA_A_R-sIPSCs. Shown are the results of experiments in 8 pairs of DLPFC L3 PNs. The statistical analysis was done with non-parametric paired Wilcoxon test (Shapiro-Wilk test, Ln-transformed data p=1×10^-4^). Bayes Factor analysis: BF_10_=4.811, median posterior d=0.906, 95% CI: [0.096, 1.849]. **D)** Top panel: examples of significant correlations between the synchGABA_A_R-sIPSC amplitudes in a DLPFC pair (left) and a PPC pair (right). Bottom panel: Violin plots comparing the Pearson’s correlation coefficients between DLPFC and PPC. Shown are the results of a Mann-Whitney U test. BF_10_=0.488, median posterior d=0.132, 95% CI: [-0.690, 1.030]. **E)** Top: Superimposed synchGABA_A_R-sIPSCs (gray traces), their average (black trace) and the double exponential curve fitted to the decay of the average (blue trace). Bottom: Violin plots comparing the decay time constants of the synchGABA_A_R-sIPSCs. Shown are the results of a Student’s t test (Shapiro Wilk test, p= 0.0621). Bayes Factor analysis: BF_10_=0.179, median posterior d=0.124, 95% CI: [0.005, 0.507].

In both PPC and DLPFC, a fraction of the total GABA_A_R-sIPSCs, here designated as synchGABA_A_R-sIPSCs, were synchronized across L3 PNs in the simultaneously recorded pairs (Fig 5A). If originating from a common presynaptic source, the synchGABA_A_R-sIPSCs should exhibit very similar timing of their onsets across cells in each pair. We found that the onsets of the synchGABA_A_R-sIPSCs differed between the L3 PNs in a pair by ≤1 ms (DLPFC: 0.45±0.16 ms, n=8 pairs; PPC: 0.56±0.12 ms, n=8 pairs), showing values highly consistent with the range of onset differences (0.1-1 ms) for IPSCs in triple recordings of a single presynaptic interneuron and two postsynaptic excitatory cells (Bartos et al., 2001). Typically, the synchGABA_A_R-sIPSCs were relatively temporally isolated (Fig 5A, Pair 1), but were also observed in bursts (Fig 5A, Pair 2). Relative to the total GABA_A_R-sIPSCs, the fraction of synchGABA_A_R-sIPSCs varied widely between cells (Fig 5B), possibly due to the high cell-to-cell variability in the total frequency of GABA_A_R-sIPSCs (Fig 2B). The voltage-gated Na^+^ channel blocker tetrodotoxin abolished the synchGABA_A_R-sIPSCs (Fig 5C), suggesting they were produced by action potential firing in interneurons presynaptic to both L3 PNs in each pair.

We reasoned that if synchGABA_A_R-sIPSCs contribute to the mechanisms synchronizing networks of L3 PNs, then the strength of the inhibitory effect from these IPSCs should be correlated across the cells in each L3 PN pair. If the inhibitory strength is not correlated, then the synchronizing effect of GABA_A_R-mediated inhibition could be less efficient or possibly absent. In most recorded pairs (14/16), the synchGABA_A_R-sIPSC amplitude was significantly correlated across L3 PNs (Fig 5D). Although the strength of the correlation varied between pairs, the correlations did not differ between DLPFC and PPC L3 PN pairs (Fig 5D). We next tested whether the synchGABA_A_R-sIPSCs have faster decay kinetics in DLPFC L3 PNs, as predicted if they contribute to the production of higher gamma oscillation frequency (Gonzalez-Burgos et al., 2015). The decay time constants of the synchGABA_A_R-sIPSCs, however, did not differ between DLPFC and PPC L3 PNs (Fig 5E), contrary to our prediction. Moreover, Bayesian analysis supported the absence of a difference in decay kinetics. These data suggest that GABA_A_R-mediated inhibition synchronizes L3 PN networks in DLPFC and PPC, but that the synchronizing effects are similar between these two areas.

### Expression levels of GABA_A_R subunit genes are similar in DLPFC and PPC L3 PNs

Previously, we reported that differences in the physiology and morphology of DLPFC and PPC L3 PNs were paralleled by patterns of differential gene expression that suggested a transcriptional basis for the different cellular phenotypes (González-Burgos et al., 2019). Consistent with the general absence of difference in GABA_A_R-sIPSCs reported here, our previous microarray study did not report any differences between DLPFC and PPC L3 PNs in the transcript levels for any GABA_A_R subunits (González-Burgos et al., 2019). However, nonsignificant differences in the expression of several subunits in combination, assessed with composite Z-scores, could still result in significant differences in GABA_A_R levels across regions. Hence, we analyzed in pools of micro-dissected L3 PNs the expression of 6 genes (Fig 6A) encoding the main alpha (GABRA1,A2, A5), beta (GABRB1, B2) and gamma (GABRG2) subunits localized in synapses, where they shape the strength and kinetics of the response to synaptically-released GABA (Lavoie et al., 1997; Engin et al., 2018; Nguyen and Nicoll, 2018). In accordance with the absence of difference in GABA_A_R-sIPSCs, we did not find any significant differences between DLPFC and PPC L3 PNs in levels of each subunit (Fig 6A) or in the composite Z-scores for all six subunits (Fig 6B).

**Figure 6.**
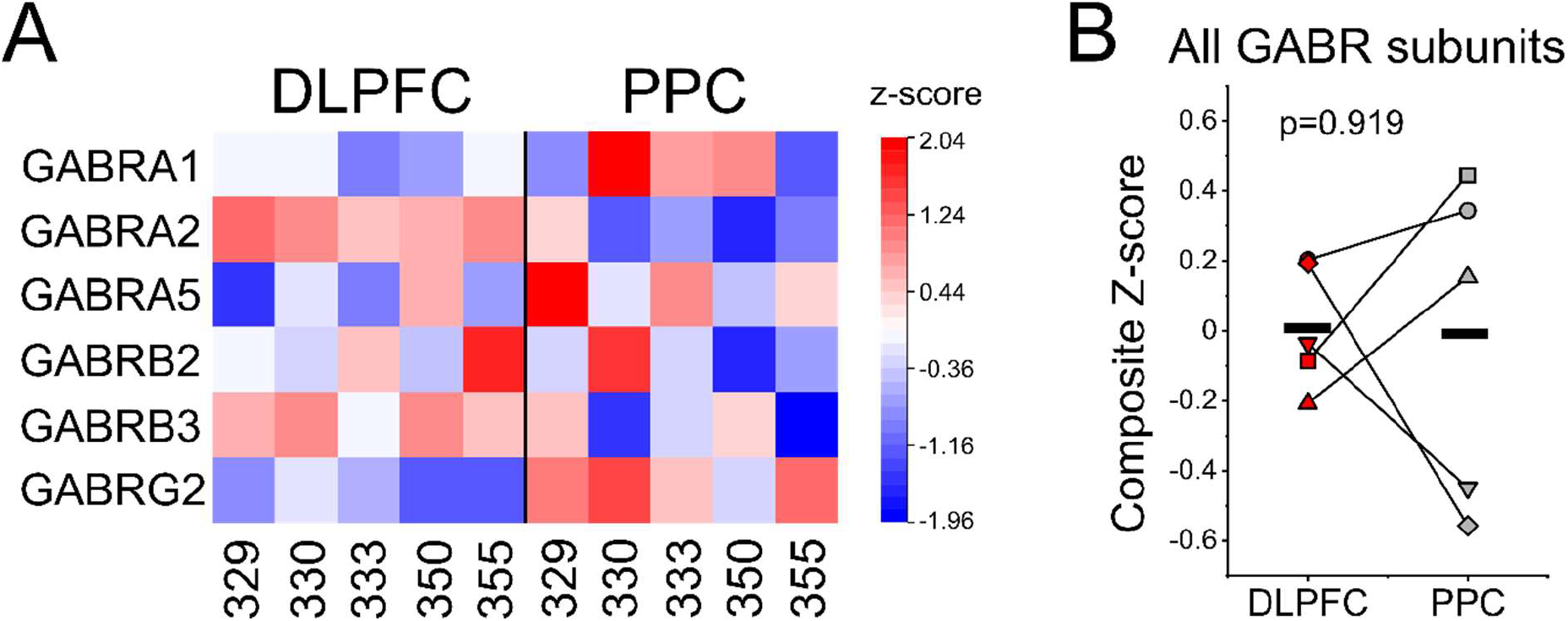
Microarray analysis of GABA_A_R gene expression in microdissected L3 PNs from DLPFC and PPC. **A)** Heat map illustrating transcript levels (expressed as Z-scores) for the 6 genes encoding the main alpha (GABRA1, A2, A5), beta (GABRB1, B2) and gamma (GABRG2) subunits known to be localized in synapses. Numbers on the X axis refer to individual macaque monkeys. **B)** Composite Z-scores generated for all the subunit transcript levels did not reveal significant GABA_A_R gene expression differences between DLPFC and PPC L3 PNs. The reported statistics is a two-tailed paired Student’s t-test (Shapiro-Wilk, p=0.660). Bayes Factor analysis: BF_10_=0.399, median posterior d=0.033, 95% CI [-0.691, 0.770]. Each animal is represented by a different symbol type, and data from a given animal are joined by a line.

### L3 PNs from DLPFC have higher dendritic spine density relative to PPC L3 PNs

Our experiments show that GABA_A_R-mediated synaptic inhibition in L3 PNs does not exhibit greater strength nor faster decay kinetics in DLPFC, arguing against the idea that the higher gamma oscillation frequency in DLPFC (Lundqvist et al., 2020) depends on unique inhibitory effects of DLPFC GABA synapses. Interestingly, L3 PNs from DLPFC exhibit higher density of dendritic spines in basal dendrites compared with L3 PNs from other cortical areas (Elston, 2007; Medalla and Luebke, 2015), and specifically from PPC (González-Burgos et al., 2019). The higher dendritic spine density suggests that L3 PNs from DLPFC could receive stronger excitatory synaptic drive, since dendritic spines are the main site of excitatory input onto PNs (DeFelipe and Farinas, 1992), and most (~97%) spines contain an excitatory synapse (Arellano et al., 2007), as verified in L3 PNs from monkey neocortex (Medalla and Luebke, 2015). By increasing network activity levels, stronger excitatory drive may determine a higher frequency of inhibition-based synchronized network oscillations without involving unique effects of inhibition (Brunel and Wang, 2003; Kirli et al., 2014).

Given that many of the L3 PNs studied here were part of our previous study quantifying dendritic spines (González-Burgos et al., 2019), we could estimate dendritic spine density (Fig 7A) in the basal dendrites of a subset of the L3 PNs in which GABA_A_R-sIPSCs were recorded in this study (DLPFC, n=14 L3 PNs; PPC, n=9 L3 PNs). Within this subset of L3 PNs, those from DLPFC had a higher density of dendritic spines (Fig 7B), and a higher ratio between basal dendrite spine density and IPSC amplitude (Fig 7C), although neither the GABA_A_R-sIPSCamplitude (Fig 7D) nor decay time constant (Fig 7E) differed between these DLPFC and PPC L3 PNs. Moreover, neither GABA_A_R-sIPSC amplitude (Pearson’s R; DLPFC: −0.547, p=0.859; PPC: −0.449, p=0.224), nor decay time constant (Pearson’s R; DLPFC: −0.004, p=0.998; PPC: 0.031, p=0.935), were correlated with dendritic spine density in DLPFC nor PPC L3 PNs.

**Figure 7.**
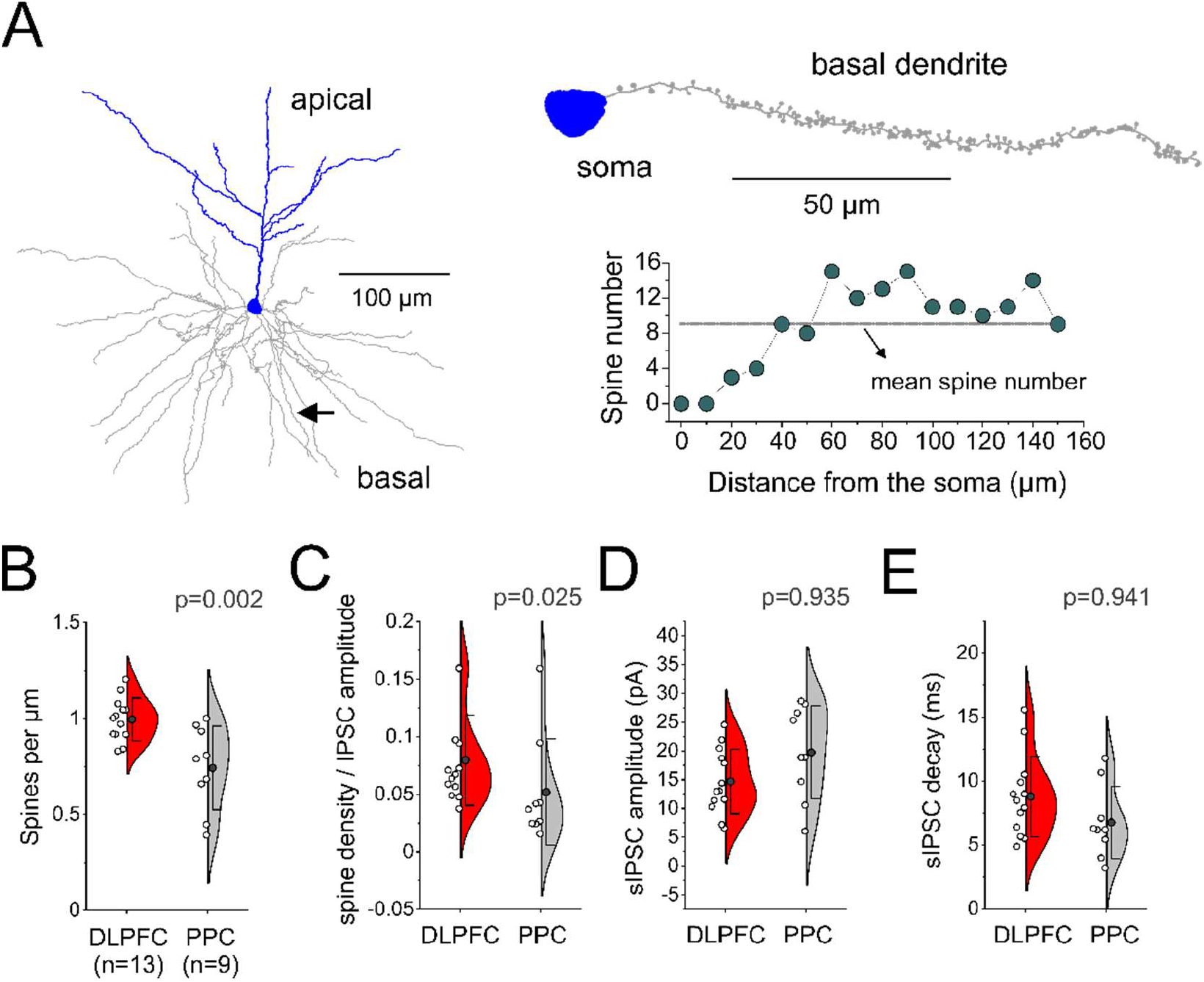
Basal dendrite dendritic spine density and GABA_A_R-mediated inhibition in L3 PNs. **A)** Left panel: reconstruction of the dendritic tree of an example L3 PN. Whereas basal dendrites were fully reconstructed, only a portion of the apical tree is shown. Arrow: single basal dendrite branch in which dendritic spines were traced. Right panel, top: the single basal dendrite branch indicated in the left panel, with the spines traced along its length. Right panel, bottom: plot of spine number as a function of distance from the soma. **B)** Spine density in basal dendrites in DLPFC and PPC L3 PNs. Comparisons performed with Student’s test (Shapiro Wilk p=0.434), Bayes Factor analysis: BF_10_=18.49, posterior median d=1.285, 95%CI: [0.321, 2.310]. **C)** Ratio between spine density and GABA_A_R-sIPSC amplitude calculated for each L3 PN (Shapiro-Wilk p=0.343 (Ln-transformed data), Bayes Factor analysis: BF_10_=2.689, posterior median d=0.796, 95% CI: [-0.014, 1.727]). **D)** GABA_A_R-sIPSC amplitude (Shapiro-Wilk p= 0.371), Bayes Factor analysis: BF_10_=0.181, posterior median d=-0.128, 95% CI: [−0.546, −0.005]) **E)** GABA_A_R-sIPSC decay time constant (Shapiro-Wilk p= 0.354), Bayes Factor analysis: BF_10_= 0.179, posterior median d=0.126, 95% CI: [0.005, 0.540]).

### Expression levels of ionotropic glutamate receptor subunit genes are higher in DLPFC than PPC L3 PNs

Given their greater number of dendritic spines, and thus excitatory synapses, it seems likely that DLPFC L3 PNs receive stronger excitatory drive, provided that, relative to PPC L3 PNs, the excitatory synapses are not weaker in DLPFC. As we did not record excitatory synaptic currents in the present study, we used data from our previous microarray analysis of gene expression (González-Burgos et al., 2019) to assess whether higher expression of ionotropic glutamate receptor genes accompanies the higher number of dendritic spines in DLPFC L3 PNs. In pools of L3 PNs micro-dissected from DLPFC and PPC, we assessed the expression of genes encoding the AMPAR (GRIA1, A2, A3, A4), and NMDAR (GRIN1A, 2A, 2B) subunits known to be localized in synapses, where they shape the strength and kinetics of the response to synaptically-released glutamate (Hansen et al., 2021; Wichmann and Kuner, 2022). For each subunit, expression levels were higher in DLPFC relative to PPC L3 PNs (Fig 8A). Moreover, composite Z-scores for all AMPAR (Fig 8B) and NMDAR (Fig 8C) subunits were significantly higher in DLPFC L3 PNs. Hence, ionotropic glutamate receptor gene expression analysis argues against the possibility that the more numerous excitatory synapses in DLPFC L3 PNs are weaker relative to those from PPC L3 PNs, supporting the idea of stronger excitatory drive in DLPFC L3 PNs.

**Figure 8.**
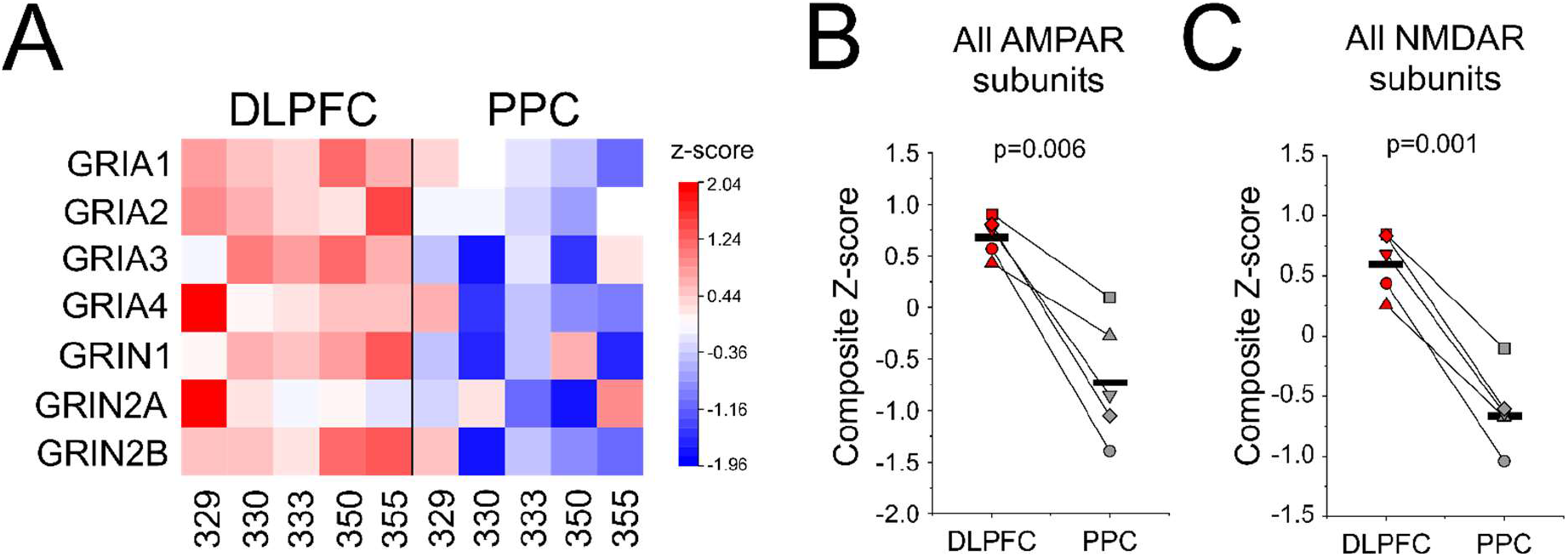
Microarray analysis of ionotropic glutamate receptors gene expression in microdissected L3 PNs from DLPFC and PPC. **A)** Heat map illustrating transcript levels (expressed as Z-scores) for 5 genes encoding the main AMPAR (GRIA1, A2, A3, A4), and NMDAR (GRIN1, 2A, 2B) subunits localized in synapses. Numbers on the X axis refer to individual macaque monkeys. **B)** Composite Z-scores generated for all the AMPAR subunit transcript levels revealed significant differences between DLPFC and PPC L3 PNs. The reported statistics is a two-tailed paired Student’s t-test (Shapiro-Wilk, p=0.178). Bayes Factor analysis: BF_10_=24.3, median posterior d=2.451, 95% CI [0.490, 4.599]. Here and in C, each animal is represented by a different symbol type, and data from a given animal are joined by a line. **B)** Composite Z-scores generated for all the NMDAR subunit transcript levels revealed significant differences between DLPFC and PPC L3 PNs. The reported statistics is a two-tailed paired Student’s t-test (Shapiro-Wilk, p=0.225). Bayes Factor analysis: BF_10_=99.5, median posterior d=3.406, 95% CI [1.017, 5.968].

### Stronger excitatory drive may generate higher gamma oscillation frequency in the DLPFC network

Our experiments showed similar strength and kinetics of synaptic inhibition in DLPFC and PPC L3 PNs and suggested that excitatory synaptic drive might be stronger in L3 PNs from DLPFC (Fig 7, 8). Previous computational modeling showed that strong external drive supports the production of high frequency (180-200 Hz) oscillations (Brunel and Wang, 2003). Here we used computational models of the local cortical microcircuit to investigate how changing the strength of excitatory synaptic drive may shape inhibition-based gamma rhythms when the properties of synaptic inhibition are held constant. Oscillations were generated, as in our previous work (Rotaru et al., 2011; Gonzalez-Burgos et al., 2015), via Pyramidal Interneuron Network Gamma (PING)-like mechanisms, in a spiking network of 80 excitatory (e) and 20 inhibitory (i) neurons coupled by recurrent synaptic connections (Fig 9A,B).

**Figure 9.**
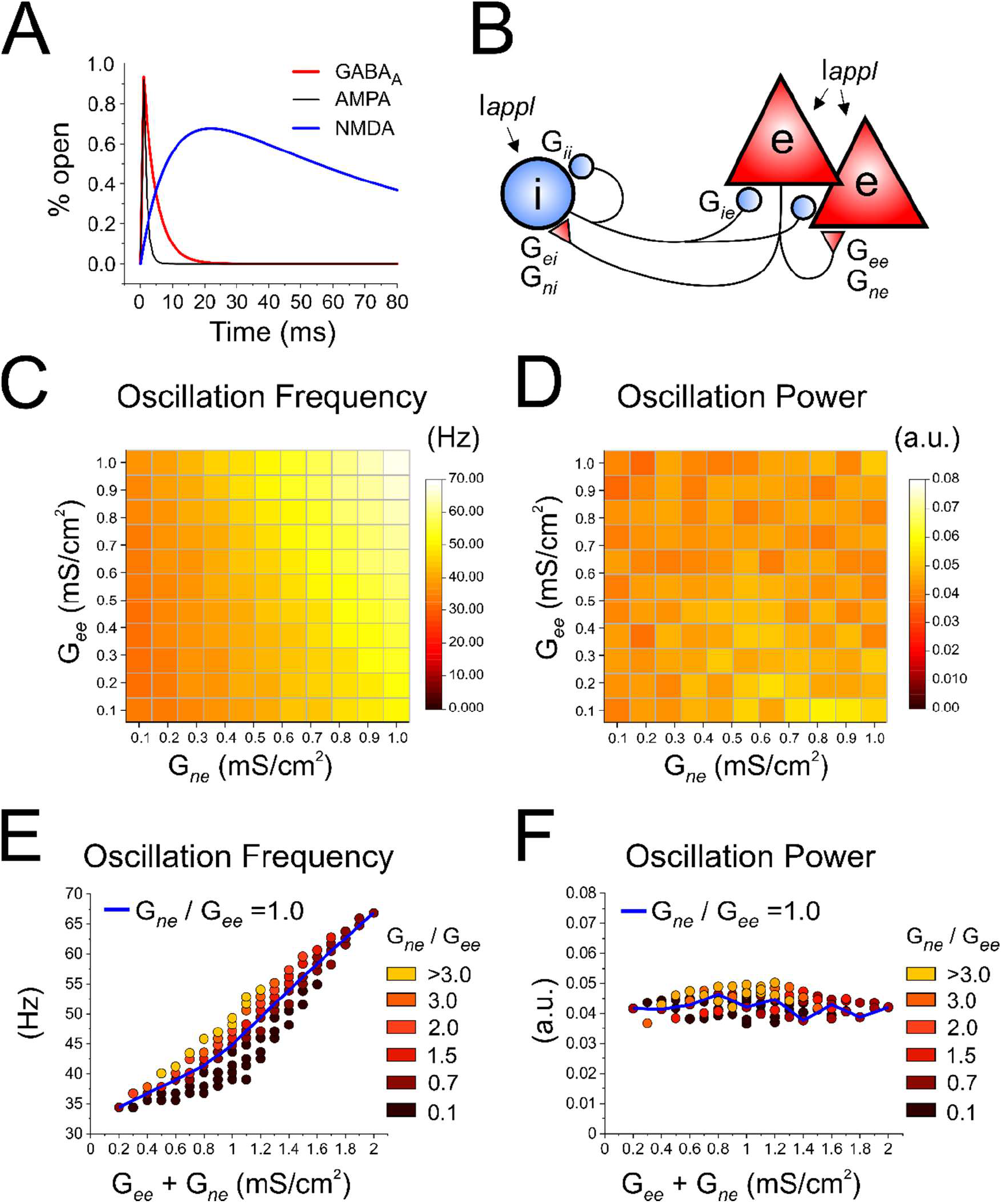
Simulations in a model of the cortical microcircuit assessing the effects of excitatory synaptic drive on gamma oscillation properties. **A)** Kinetic properties of the excitatory AMPA (G_*ee*_ and G_*ei*_) and NMDA (G_*ne*_ and G_*ni*_) conductance, and the inhibitory GABA_A_ (G_*ie*_ and G_*ii*_) conductance implemented at the synaptic connections in the model network. **B)** Schematic representation of the network connectivity. *l_appl_* is the excitatory current drive directly applied to the neurons independent of synaptic inputs. **C)** Heatmap showing the peak oscillation frequency for networks with different values of AMPA and NMDA conductance at the recurrent connections (G_*ee*_ and G_*ne*_). **D)** Heatmap showing the peak oscillation frequency for networks with different values of G_*ee*_ and G_*ne*_. **E)** Scatter plot of oscillation frequency as a function of total excitatory drive (G_*ee*_+G_*ne*_) for the different NMDA/AMPA ratios (G_*ne*_/G_*ee*_). **F)** Scatter plot of oscillation power as a function of G_*ee*_+G_*ne*_ for different G_*ne*_/G_*ee*_ ratios.

Although we did not record excitatory synaptic currents for this study, in cortical PNs these currents are typically mediated via both AMPAR- and NMDAR-mediated components, as in L3 PNs from monkey DLPFC (Gonzalez-Burgos et al., 2008). Hence, the recurrent excitatory synapses onto the e cells in the network were modeled as having a combination of AMPAR-mediated (G_*ee*_) and NMDAR-mediated (G_*ne*_) excitatory synaptic conductances, simultaneously gated when each input is activated (Fig 9A,B). The external excitatory current applied onto the e cells and i cells (*l_appl_* in Fig 9B) was held constant, to focus on changes in the recurrent excitatory synaptic connections.

We simulated oscillations in different networks, each exhibiting a different level of total excitatory synaptic drive (G_*ee*_+G_*ne*_). To vary the excitatory synaptic drive from network to network, first, we ramped-up G_*ee*_+G_*ne*_ by incrementing G_*ee*_ and *G_*ne*_* in identical amounts, thus keeping the NMDAR/AMPAR ratio constant (G_*ne*_/G_*ee*_ = 1). Next, we increased G_*ee*_+G_*ne*_ by augmenting G_*ee*_ and G_*ne*_ in proportions that increased (G_*ne*_/G_*ee*_ > 1) or decreased (G_*ne*_/G_*ee*_< 1) the NMDAR/AMPAR ratio. To mimic the absence of difference in synaptic inhibition between DLPFC and PPC observed in the experiments, we held constant the GABA_A_R-mediated inhibitory conductance onto e (G_*ie*_) and i (G_*ii*_) cells, and the AMPAR-mediated (G_*ei*_) and NMDAR-mediated (G_*ni*_) excitatory conductance onto the i neurons (Fig 9A,B). As we parametrically varied G_*ee*_ and G_*ne*_, the simulations were performed in 100 independent networks.

We found that networks with stronger excitatory drive (G_*ee*_+G_*ne*_) produced higher peak oscillation frequency (Fig 9C), without showing consistent differences in the peak oscillation power (Fig 9D). Moreover, the differences in oscillation frequency between networks were mainly due to a stronger total excitatory drive G_*ee*_+G_*ne*_, since increasing only G_*ee*_ or G_*ne*_, while keeping low levels of the other conductance, produced relatively minor changes in oscillation properties, as depicted by the heatmaps in Fig 9C,D. The heatmaps also suggested that increasing G_*ne*_ had some greater effects on the oscillation frequency than increasing G_*ee*_ (Fig 9C). In order to assess the role of G_*ee*_ and G_*ne*_ in greater detail, we built scatter plots of peak oscillation frequency (Fig 9E) and power (Fig 9F) as a function of G_*ee*_+G_*ne*_, for the different values of G_*ne*_/G_*ee*_. These plots showed that the oscillation frequency significantly increased from network to network as we increased G_*ee*_+G_*ne*_, largely along the line of G_*ne*_/G_*ee*_ = 1 (Fig 9E). Moreover, the increase in frequency with G_*ee*_+G_*ne*_, was observed in all networks irrespective of the G_*ne*_/G_*ee*_ value for each network (Fig 9E), consistent with the idea that the frequency of the oscillations increased mostly by effect of the increase in total excitatory drive. Interestingly, for most G_*ee*_+G_*ne*_ values, the oscillation frequency was lower in networks with G_*ne*_/G_*ee*_ < 1 and higher in networks with G_*ne*_/G_*ee*_ > 1 (Fig 9E). The oscillation power, in contrast, did not differ consistently between networks with different values of G_*ee*_+G_*ne*_ or G_*ne*_/G_*ee*_ (Fig 9F). Our data therefore suggest that the strength of excitatory synaptic drive from recurrent excitation is a crucial determinant of gamma oscillation frequency. Moreover, the results of our simulations suggest that if a network exhibits higher NMDAR/AMPAR ratio at the recurrent excitatory synapses, then generating gamma oscillations of higher frequency is facilitated.

## Discussion

We investigated L3 PN properties that might contribute to the production of a higher frequency of gamma oscillations in primate DLPFC relative to PPC. Given the crucial role of GABA_A_R-mediated inhibition in the generation of synchronized gamma oscillations, we first studied GABA_A_R-sIPSCs. We found evidence that GABA_A_R-mediated inhibition synchronizes L3 PNs in DLPFC and PPC but found that neither the strength nor kinetics of GABA_A_R-sIPSCs differed between these areas. Likewise, GABA_A_R gene expression in L3 PNs did not show any DLPFC-PPC differences. Interestingly, in the absence of differences in GABA_A_R-sIPSCs, DLPFC L3 PNs exhibited higher number of dendritic spines and higher expression of AMPAR and NMDAR genes, supporting the idea of stronger excitatory synaptic drive in DLPFC L3 PNs. Simulations of inhibition-based gamma oscillations showed that a stronger recurrent excitatory drive substantially increased the network oscillation frequency. Hence, the mechanisms producing higher frequency gamma oscillations in DLPFC may involve stronger recurrent excitatory drive.

### GABA_A_R-mediated IPSCs show similar properties in DLPFC and PPC layer 3 pyramidal cells

Previous studies demonstrated that gamma oscillation frequency increases across the primate cortex hierarchy, as it is similar in the low-order areas V1 and V2 (Roberts et al., 2013), but is higher in V4, a higher-order region of the monkey cortex (Rols et al., 2001; Tallon-Baudry, 2009). Moreover, oscillations elicited with transcranial magnetic stimulation have higher frequency in frontal than parietal or V1 areas of the human cortex (Rosanova et al., 2009; Ferrarelli et al., 2012). In addition, during a working memory task, gamma oscillation frequency increases from V4 towards higher-order areas PPC and DLPFC, and is higher in DLPFC relative to PPC (Lundqvist et al., 2020). In our study, neither GABA_A_R-sIPSC amplitude nor decay kinetics differed between DLPFC and PPC, including the synchGABA_A_R-sIPSCs presumably involved in inhibition-based synchrony. Moreover, a previous study showed that GABA_A_R-sIPSC amplitude and decay kinetics do not differ between L3 PNs from monkey DLPFC and V1 (Amatrudo et al., 2012). Hence, the available data argue against the idea that differences in GABA_A_R-mediated inhibition generate different gamma oscillation properties in different areas of the primate neocortex.

Our results additionally show that both DLPFC and PPC L3 PNs display a wide range of GABA_A_R-sIPSC decay time constant values (<5 ms to >20 ms). In our previous work, L3 PN networks with IPSC decay time constants varying within that range produced oscillation frequencies spanning from beta (~20 Hz) to gamma (~55 Hz) bands (Gonzalez-Burgos et al., 2015). These wide distributions of IPSC decay time constants may therefore provide a menu for selectively recruiting IPSCs with faster or slower kinetics to generate gamma oscillations with different frequency.

The mechanisms by which subsets of GABA_A_R-sIPSCs with different decay kinetics could be selectively recruited are, however, currently unclear. For example, all DLPFC and PPC L3 PNs showed wide distributions of IPSC decay constants, suggesting that GABA_A_R-IPSCs with faster or slower decay kinetics are not associated with specific L3 PN subtypes. Separate subsets of IPSCs are selectively recruited by activation of interneurons of different subtypes. However, in the rodent neocortex, presynaptic interneurons of different subtype elicit in postsynaptic pyramidal cells GABA_A_R-IPSCs with similar decay kinetics (Xiang et al., 2002; Ali and Thomson, 2008; Galarreta et al., 2008; Donato et al., 2021; Campagnola et al., 2022). Likewise, different subtypes of presynaptic interneurons elicit GABA_A_R-IPSCs with similar decay kinetics in L3 PNs from monkey DLPFC (Gonzalez-Burgos et al., 2005).

### Excitatory synaptic drive may control oscillation frequency in cortical networks with similar GABA_A_R-mediated inhibition

The absence of difference in GABA_A_R-sIPSC properties suggests that the different gamma oscillation frequency in DLPFC and PPC (Lundqvist et al., 2020) does not involve different effects of inhibition. Here we propose that the higher gamma oscillation frequency in DLPFC might be associated with stronger recurrent excitatory drive. This postulate is supported by findings showing that L3 PNs have higher dendritic spine density in DLPFC relative to PPC (Elston, 2000; Elston et al., 2001; Elston, 2007), and specifically in DLPFC area 46 relative to PPC areas 7a/LIP (González-Burgos et al., 2019). Here, we show in a sample of DLPFC and PPC L3 PNs that, despite showing similar GABA_A_R-sIPSC properties, those from DLPFC have higher spine density in basal dendrites. These results support the idea that excitatory drive onto DLPFC L3 PNs is stronger, unless the more numerous excitatory synapses in DLPFC are weaker. Arguing against weaker glutamate synapses in DLPFC, we found higher levels of AMPAR and NMDAR subunit transcripts in DLPFC than PPC L3 PNs.

Basal dendrites, where DLPFC L3 PNs show higher number of spines (González-Burgos et al., 2019), are the primary target of recurrent excitatory connections (Markram et al., 1997; Gökçe et al., 2016), and display specialized NMDAR-mediated mechanisms that could boost recurrent excitation (Schiller et al., 2000; Nevian et al., 2007; Major et al., 2008). We found higher levels of NMDAR subunit transcripts in DLPFC L3 PNs, including for the GluN2B subunits which may facilitate recurrent excitation (Wang and Arnsten, 2015; Wang, 2020), hence suggesting greater NMDAR-mediated boosting of recurrent excitation in DLPFC.

We assessed the role of recurrent excitatory drive simulating inhibition-based rhythms as we varied between different networks the AMPAR- and NMDAR-mediated contribution to recurrent excitation. The simulations showed that the oscillation frequency was higher in networks with greater total excitatory drive (G_*ee*_+G_*ne*_), and that a higher NMDAR/AMPAR ratio at the recurrent synapses between excitatory cells could contribute to higher oscillation frequency. Hence, in the absence of differences in synaptic inhibition, stronger recurrent excitation may contribute to the higher oscillation frequency in higher-order cortical areas (Rols et al., 2001; Rosanova et al., 2009; Tallon-Baudry, 2009; Ferrarelli et al., 2012; Lundqvist et al., 2020).

Importantly, we found that DLPFC L3 PNs exhibit higher levels of transcripts for both NMDAR and AMPAR subunits. Since DLPFC neurons have larger dendrites and higher dendritic spine density relative to PPC L3 PNs (González-Burgos et al., 2019), the higher expression of ionotropic glutamate receptor genes could primarily reflect the higher synapse number. Future studies quantifying receptors in the postsynaptic membrane or recording excitatory synaptic currents are therefore necessary to determine whether the NMDAR/AMPAR ratio differs between DLPFC and PPC L3 PNs.

### Functional implications

Simulations based on our experimental results suggested that, without differences in GABA_A_R-mediated synaptic inhibition, gamma oscillation frequency is regulated by recurrent excitatory drive. The mechanisms revealed by our simulations may apply to separate networks that, such as the DLPFC and PPC, differ in the total number of synapses per excitatory neuron. But we simulated differences in excitatory drive without distinguishing, hence being independent of, whether stronger excitatory drive was due to more numerous or stronger excitatory synapses. Consequently, similar effects of excitatory drive may be expected for distinct states of a single network, having larger or smaller fractions of synapses recruited out of the total excitatory synapses available. Our results are therefore consistent with the idea that oscillation frequency might be dynamically regulated with stable microcircuit structure within a cortical region, by adjusting the amount of excitation recruited. Consistent with dynamic modulation, in macaque cortex areas V1 or V2, the oscillation frequency switches reversibly between beta (~20 Hz) and gamma (~50 Hz) bands, as a function of visual stimulus contrast (Ray and Maunsell, 2010; Roberts et al., 2013). It remains to be determined the extent to which differences in microcircuit structure and dynamic modulation of excitatory drive contribute to the different oscillation frequency in DLPFC and PPC during working memory tasks.

While supporting the concept that stronger excitatory drive may be a mechanism to generate higher oscillation frequency in the DLPFC network, our simulations did not attempt to exactly reproduce the oscillation properties observed during working memory tasks (Lundqvist et al., 2020). Building on our proof-of-concept tests, however, future studies could determine the parameters necessary to mimic the oscillation properties observed in the intact rhesus macaque cortex throughout different phases of a working memory task, particularly during the delay period in the absence of cue-driven inputs.

Although we simulated only the effects of local excitatory drive, we speculate that in the intact cortex the frequency of inhibition-based oscillations may be determined by interactions between local and external excitatory drive. Specifically, our results suggest that with external drive of similar strength, which we mimicked by a constant I_*appl*_, the oscillation frequency might be higher in networks that, such as the DLPFC, may exhibit stronger local recurrent excitation. Interestingly, simultaneous recording from DLPFC and PPC neurons showed significant long-range reciprocal excitation with similar strength in the DLPFC-to-PPC and PPC-to-DLPFC directions during the delay period of working memory tasks (Hart and Huk, 2020). Moreover, inactivation experiments with reversible cooling showed that, during working memory tasks, external input flowing to DLPFC from PPC or in the reverse direction, both are predominant sources of excitatory drive for neurons located in each of these two areas (Chafee and Goldman-Rakic, 2000). Hence, external excitatory input of similar strength flowing bidirectionally between DLPFC and PPC, together with stronger local recurrent excitation in DLPFC, may contribute to the production of higher gamma oscillation frequency in DLPFC.

## Acknowledgements

We thank Kate Gurnsey for assistance in surgical procedures and Olga Krimer for excellent assistance in neuronal reconstructions.

## References

Ali AB, Thomson AM (2008) Synaptic alpha 5 subunit-containing GABAA receptors mediate IPSPs elicited by dendrite-preferring cells in rat neocortex. Cereb Cortex 18:1260–1271.

Amatrudo JM, Weaver CM, Crimins JL, Hof PR, Rosene DL, Luebke JI (2012) Influence of highly distinctive structural properties on the excitability of pyramidal neurons in monkey visual and prefrontal cortices. J Neurosci 32:13644–13660.

Arellano JI, Espinosa A, Fairen A, Yuste R, DeFelipe J (2007) Non-synaptic dendritic spines in neocortex. Neuroscience 145:464–469.

Banaie Boroujeni K, Tiesinga P, Womelsdorf T (2021) Interneuron-specific gamma synchronization indexes cue uncertainty and prediction errors in lateral prefrontal and anterior cingulate cortex. Elife 10.

Bartos M, Vida I, Frotscher M, Geiger JR, Jonas P (2001) Rapid signaling at inhibitory synapses in a dentate gyrus interneuron network. J Neurosci 21:2687–2698.

Bastos AM, Loonis R, Kornblith S, Lundqvist M, Miller EK (2018) Laminar recordings in frontal cortex suggest distinct layers for maintenance and control of working memory. Proc Natl Acad Sci U S A 115:1117–1122.

Brunel N, Wang XJ (2003) What determines the frequency of fast network oscillations with irregular neural discharges? I. Synaptic dynamics and excitation-inhibition balance. J Neurophysiol 90:415–430.

Buzsaki G, Wang XJ (2012) Mechanisms of gamma oscillations. Annu Rev Neurosci 35:203–225.

Campagnola L et al. (2022) Local connectivity and synaptic dynamics in mouse and human neocortex. Science 375:eabj5861.

Chafee MV, Goldman-Rakic PS (2000) Inactivation of parietal and prefrontal cortex reveals interdependence of neural activity during memory-guided saccades. J Neurophysiol 83:1550–1566.

Christophel TB, Klink PC, Spitzer B, Roelfsema PR, Haynes JD (2017) The Distributed Nature of Working Memory. Trends Cogn Sci 21:111–124.

Datta D, Arion D, Lewis DA (2015) Developmental Expression Patterns of GABAA Receptor Subunits in Layer 3 and 5 Pyramidal Cells of Monkey Prefrontal Cortex. Cereb Cortex 25:2295–2305.

DeFelipe J, Farinas I (1992) The pyramidal neuron of the cerebral cortex: morphological and chemical characteristics of the synaptic inputs. Progress in Neurobiology 39:563–607.

Donato C, Cabezas C, Aguirre A, Lourenco J, Poiter M-C, Zorilla de San Martin J, Bacci A (2021) Molecular signature and target-specificity of inhibitory circuits formed by Martinotti cells in the mouse barrel cortex. bioRxiv https://doi.org/10.1101/2021.08.16.455953

Elston GN (2000) Pyramidal cells of the frontal lobe: all the more spinous to think with. J Neurosci 20:RC95.

Elston GN (2007) Specialization of the Neocortical Pyramidal Cell during Primate Evolution. In: Evolution of Nervous Systems (Kaas JH, ed): Academic Press.

Elston GN, Benavides-Piccione R, DeFelipe J (2001) The pyramidal cell in cognition: a comparative study in human and monkey. J Neurosci 21:RC163.

Engin E, Benham RS, Rudolph U (2018) An Emerging Circuit Pharmacology of GABA(A) Receptors. Trends Pharmacol Sci 39:710–732.

Ferrarelli F, Sarasso S, Guller Y, Riedner BA, Peterson MJ, Bellesi M, Massimini M, Postle BR, Tononi G (2012) Reduced natural oscillatory frequency of frontal thalamocortical circuits in schizophrenia. Arch Gen Psychiatry 69:766–774.

Galarreta M, Erdelyi F, Szabo G, Hestrin S (2008) Cannabinoid sensitivity and synaptic properties of 2 GABAergic networks in the neocortex. Cereb Cortex 18:2296–2305.

Gökçe O, Bonhoeffer T, Scheuss V (2016) Clusters of synaptic inputs on dendrites of layer 5 pyramidal cells in mouse visual cortex. Elife 5.

Goldman-Rakic PS (1988) Topography of cognition: parallel distributed networks in primate association cortex. Annu Rev Neurosci 11:137–156.

Gonzalez-Burgos G, Krimer LS, Povysheva NV, Barrionuevo G, Lewis DA (2005) Functional properties of fast spiking interneurons and their synaptic connections with pyramidal cells in primate dorsolateral prefrontal cortex. Journal of Neurophysiology 93:942–953.

Gonzalez-Burgos G, Kroener S, Zaitsev AV, Povysheva NV, Krimer LS, Barrionuevo G, Lewis DA (2008) Functional maturation of excitatory synapses in layer 3 pyramidal neurons during postnatal development of the primate prefrontal cortex. Cereb Cortex 18:626–637.

Gonzalez-Burgos G, Miyamae T, Pafundo DE, Yoshino H, Rotaru DC, Hoftman G, Datta D, Zhang Y, Hammond M, Sampson AR, Fish KN, Ermentrout GB, ewis DA (2015) Functional Maturation of GABA Synapses During Postnatal Development of the Monkey Dorsolateral Prefrontal Cortex. Cereb Cortex 25:4076–4093.

González-Burgos G, Miyamae T, Krimer Y, Gulchina Y, Pafundo DE, Krimer O, Bazmi H, Arion D, Enwright JF, Fish KN, Lewis DA (2019) Distinct Properties of Layer 3 Pyramidal Neurons from Prefrontal and Parietal Areas of the Monkey Neocortex. J Neurosci 39:7277–7290.

Hansen KB, Wollmuth LP, Bowie D, Furukawa H, Menniti FS, Sobolevsky AI, Swanson GT, Swanger SA, Greger IH, Nakagawa T, McBain CJ, Jayaraman V, Low CM, Dell’Acqua ML, Diamond JS, Camp CR, Perszyk RE, Yuan H, Traynelis SF (2021) Structure, Function, and Pharmacology of Glutamate Receptor Ion Channels. Pharmacol Rev 73:298–487.

Hart E, Huk AC (2020) Recurrent circuit dynamics underlie persistent activity in the macaque frontoparietal network. Elife 9:e52460.

Howard MW, Rizzu DS, Caplan JB, Madsen JR, Lisman J, Aschenbrenner-Scheiber R, Schulze-Bonhage A, Kahana MJ (2003) Gamma oscillations correlate with working memory load in humans. Cerebral Cortex 13:1369–1374.

Irizarry RA, Hobbs B, Collin F, Beazer-Barclay YD, Antonellis KJ, Scherf U, Speed TP (2003) Exploration, normalization, and summaries of high density oligonucleotide array probe level data. Biostatistics 4:249–264.

Keysers C, Gazzola V, Wagenmakers EJ (2020) Using Bayes factor hypothesis testing in neuroscience to establish evidence of absence. Nat Neurosci 23:788–799.

Kirli KK, Ermentrout GB, Cho RY (2014) Computational study of NMDA conductance and cortical oscillations in schizophrenia. Front Comput Neurosci 8:133.

Kudoh SN, Taguchi T (2002) A simple exploratory algorithm for the accurate and fast detection of spontaneous synaptic events. Biosens Bioelectron 17:773–782.

Lavoie AM, Tingey JJ, Harrison NL, Pritchett DB, Twyman RE (1997) Activation and deactivation rates of recombinant GABA(A) receptor channels are dependent on alpha-subunit isoform. Biophys J 73:2518–2526.

Leavitt ML, Mendoza-Halliday D, Martinez-Trujillo JC (2017) Sustained Activity Encoding Working Memories: Not Fully Distributed. Trends Neurosci 40:328–346.

Lundqvist M, Bastos AM, Miller EK (2020) Preservation and Changes in Oscillatory Dynamics across the Cortical Hierarchy. J Cogn Neurosci 32:2024–2035.

Lundqvist M, Herman P, Palva M, Palva S, Silverstein D, Lansner A (2013) Stimulus detection rate and latency, firing rates and 1-40Hz oscillatory power are modulated by infra-slow fluctuations in a bistable attractor network model. Neuroimage 83:458–471.

Lundqvist M, Rose J, Herman P, Brincat SL, Buschman TJ, Miller EK (2016) Gamma and Beta Bursts Underlie Working Memory. Neuron 90:152–164.

Major G, Polsky A, Denk W, Schiller J, Tank DW (2008) Spatiotemporally graded NMDA spike/plateau potentials in basal dendrites of neocortical pyramidal neurons. J Neurophysiol 99:2584–2601.

Markov NT et al. (2014) A weighted and directed interareal connectivity matrix for macaque cerebral cortex. Cereb Cortex 24:17–36.

Markram H, Lubke J, Frotscher M, Roth A, Sakmann B (1997) Physiology and anatomy of synaptic connections between thick tufted pyramidal neurones in the developing rat neocortex. Journal of Physiology 500:409–440.

Medalla M, Luebke JI (2015) Diversity of glutamatergic synaptic strength in lateral prefrontal versus primary visual cortices in the rhesus monkey. J Neurosci 35:112–127.

Medalla M, Gilman JP, Wang JY, Luebke JI (2017) Strength and Diversity of Inhibitory Signaling Differentiates Primate Anterior Cingulate from Lateral Prefrontal Cortex. J Neurosci 37:4717–4734.

Miller EK, Lundqvist M, Bastos AM (2018) Working Memory 2.0. Neuron 100:463–475.

Miyamae T, Chen K, Lewis DA, Gonzalez Burgos G (2017) Distinct physiological maturation of parvalbumin-positive neuron subtypes in mouse prefrontal cortex Journal of Neuroscience 37:4883–4902.

Nevian T, Larkum ME, Polsky A, Schiller J (2007) Properties of basal dendrites of layer 5 pyramidal neurons: a direct patch-clamp recording study. Nat Neurosci 10:206–214.

Nguyen QA, Nicoll RA (2018) The GABA(A) Receptor β Subunit Is Required for Inhibitory Transmission. Neuron 98:718–725.e713.

Pesaran B, Pezaris JS, Sahani M, Mitra PP, Andersen RA (2002) Temporal structure in neuronal activity during working memory in macaque parietal cortex. Nat Neurosci 5:805–811.

Ray S, Maunsell JH (2010) Differences in gamma frequencies across visual cortex restrict their possible use in computation. Neuron 67:885–896.

Roberts MJ, Lowet E, Brunet NM, Ter WM, Tiesinga P, Fries P, De Weerd P (2013) Robust Gamma Coherence between Macaque V1 and V2 by Dynamic Frequency Matching. Neuron 78:523–536.

Rols G, Tallon-Baudry C, Girard P, Bertrand O, Bullier J (2001) Cortical mapping of gamma oscillations in areas V1 and V4 of the macaque monkey. Vis Neurosci 18:527–540.

Rosanova M, Casali A, Bellina V, Resta F, Mariotti M, Massimini M (2009) Natural frequencies of human corticothalamic circuits. J Neurosci 29:7679–7685.

Rotaru DC, Yoshino H, Lewis DA, Ermentrout GB, Gonzalez-Burgos G (2011) Glutamate receptor subtypes mediating synaptic activation of prefrontal cortex neurons: relevance for schizophrenia. J Neurosci 31:142–156.

Rothman JS, Silver RA (2018) NeuroMatic: An Integrated Open-Source Software Toolkit for Acquisition, Analysis and Simulation of Electrophysiological Data. Front Neuroinform 12:14.

Roux F, Wibral M, Mohr HM, Singer W, Uhlhaas PJ (2012) Gamma-band activity in human prefrontal cortex codes for the number of relevant items maintained in working memory. J Neurosci 32:12411–12420.

Rudolph U, Knoflach F (2011) Beyond classical benzodiazepines: novel therapeutic potential of GABAA receptor subtypes. Nat Rev Drug Discov 10:685–697.

Schiller J, Major G, Koester HJ, Schiller Y (2000) NMDA spikes in basal dendrites of cortical pyramidal neurons. Nature 404:285–289.

Selemon LD, Goldman-Rakic PS (1988) Common cortical and subcortical targets of the dorsolateral prefrontal and posterior parietal cortices in the rhesus monkey: evidence for a distributed neural network subserving spatially guided behavior. J Neurosci 8:4049–4068.

Tallon-Baudry C (2009) The roles of gamma-band oscillatory synchrony in human visual cognition. Frontiers in Biosciences 14:321–332.

Todd JJ, Marois R (2005) Posterior parietal cortex activity predicts individual differences in visual short-term memory capacity. Cogn Affect Behav Neurosci 5:144–155.

van Doorn J, van den Bergh D, Böhm U, Dablander F, Derks K, Draws T, Etz A, Evans NJ, Gronau QF, Haaf JM, Hinne M, Kucharský Š, Ly A, Marsman M, Matzke D, Gupta A, Sarafoglou A, Stefan A, Voelkel JG, Wagenmakers EJ (2021) The JASP guidelines for conducting and reporting a Bayesian analysis. Psychon Bull Rev 28:813–826.

Wang M, Arnsten AF (2015) Contribution of NMDA receptors to dorsolateral prefrontal cortical networks in primates. Neurosci Bull 31:191–197.

Wang XJ (2020) Macroscopic gradients of synaptic excitation and inhibition in the neocortex. Nat Rev Neurosci 21:169–178.

Wang Z, Singh B, Zhou X, Constantinidis C (2022) Strong gamma frequency oscillations in the adolescent prefrontal cortex. J Neurosci 42:2917–2929.

Whittington MA, Traub RD, Kopell N, Ermentrout B, Buhl EH (2000) Inhibition-based rhythms: experimental and mathematical observations on network dynamics. Int J Psychophysiol 38:315–336.

Wichmann C, Kuner T (2022) Heterogeneity of glutamatergic synapses: cellular mechanisms and network consequences. Physiol Rev 102:269–318.

Xiang Z, Huguenard JR, Prince DA (2002) Synaptic inhibition of pyramidal cells evoked by different interneuronal subtypes in layer v of rat visual cortex. J Neurophysiol 88:740–750.

Yu Q, Panichello MF, Cai Y, Postle BR, Buschman TJ (2020) Delay-period activity in frontal, parietal, and occipital cortex tracks noise and biases in visual working memory. PLoS Biol 18:e3000854.

